# Engineering erythrocytes into synthetic cells using cell-free gene expression

**DOI:** 10.64898/2026.08.20.745953

**Authors:** David Garenne, Seth Thompson, Vincent Noireaux

**Author notes:** **Corresponding author:** Vincent Noireaux.

## Abstract

Erythrocytes, commonly known as red blood cells (RBCs), constitute the most abundant cell type in vertebrate mammals. Due to their unique biological and physical attributes, RBCs have been the focus of extensive research in biomedical engineering. Methods have been developed to transform RBCs into adaptable carriers for molecular payloads, thereby extending their functional capabilities beyond what they naturally transport and accomplish. Concurrently with RBC’s applied science, cell-free gene expression (CFE) has advanced into a tractable technology that can be integrated with a broad range of materials. In this work, we harness the advantages of CFE to engineer RBCs into hybrid synthetic cells. We encapsulate CFE reactions within RBC ghosts to execute elementary gene circuits, including biosensors, and to synthesize phages from their genomes. Furthermore, we engineer and functionalize the outer membrane of mature RBCs to attach diverse payloads, such as a SARS-CoV-2 antigen recognized by a specific antibody. CFE interfaces remarkably well with RBCs, enabling their rapid, low-cost transformation into red blood synthetic cells (RBSCs) with potential biomedical and biotechnological applications.

## Introduction

Cell-free gene expression (CFE), also known as cell-free transcription-translation (TXTL), enables DNA-driven protein synthesis in just a few hours, across reaction scales spanning eighteen orders of magnitude, from femtoliters to cubic meters [1–4]. Over the past couple of decades, CFE has been transformed into an effective and accessible technology with an ever-growing array of applications, most notably in bioengineering [5–13]. With protein synthesis yields reaching several mg/ml, CFE delivers sufficient product for applications as diverse as protein screening and biomanufacturing [14–22], prototyping gene parts and circuits with a design-build-test-learn cycle (DBTLC) that outcompetes other methods [23–32], and creating biosensors [6,33–37]. CFE was also developed for integration with a wide variety of materials, extending beyond ordinary laboratory plasticwares, including, among others, encapsulation into cell-sized liposomes to build bottom-up synthetic cells [38–47], incorporation into solid-state chips [48–54], and freeze-drying onto paper substrates [55–58]. Some areas of CFE with potential applications, however, remain underexplored. One of such uncharted territories is the engineering of cells. The motivation to pursue the research presented in this manuscript began by asking what type of living cells could be engineered using CFE, and to what extent they could be engineered for biomedical and biotechnological applications. Due to their distinctive biological and physical attributes, erythrocytes, commonly known as red blood cells (RBCs), emerged as an optimal cell type for evaluating CFE-mediated cell engineering.

Erythrocytes (RBCs) are ubiquitous in mammals and typically constitute the most abundant cell type in these organisms. RBCs constitute more than 80% of the number of human cells [59]. Mature RBCs from vertebrate mammals lack nuclei and genetic information, and do not contain functional nucleases. They do not divide and have a lifetime of 50-120 days before being recycled in the spleen and liver. They can be readily isolated from blood and safely administered intravenously back into the circulatory system. With an average diameter of 7-8 µm, RBCs are relatively small, physically flexible, and naturally circulate everywhere in vertebrate mammals via the circulatory system. With such biological and physical attributes, RBCs are uniquely designed to deliver payloads to the body by acting as circulatory vehicles. They have been the focus of bioengineering geared towards biomedical applications [60]. RBCs have longer lifetimes in blood circulation than any other drug carriers so far engineered [61]. Diverse payloads have been encapsulated into RBCs and RBC ghosts, including small drug molecules, proteins and antibodies, nucleic acids, and nanoparticles [62–67]. Several methods have been devised for the intracellular encapsulation of biologics into RBCs, with hypotonic hemolysis being the most common one. When placed in a hypotonic solution, the membrane of RBCs forms transient pores of 20-50 nm diameter, allowing molecules to enter. The membrane reseals once the RBCs are placed in an isotonic solution. Other methods, including electroporation and extrusion, have been used to encapsulate highly charged or large molecular payloads [62,63]. Several methods have also been developed for extracellular loading onto the outer membrane of RBCs [62,63], which does not require damaging the RBCs’ internal structure. Molecules can be covalently linked to RBC’s surface proteins using crosslinkers, high-affinity systems like biotin-streptavidin or specific binding peptides [68–71]. Nanocarriers can be non-covalently adsorbed onto the RBC’s surface via electrostatic or hydrophobic interactions [72–75]. Electro insertion has also been shown to work [76]. Additionally, toxin and chemical sequestration has been demonstrated using RBCs as a realization of nano-sponges [77–81]. Advanced approaches, genetic engineering especially, have also been demonstrated but are technically more demanding. Hematopoietic stem cells must be genetically modified to express therapeutic proteins within or on the surface of the resulting mature RBCs [82,83].

In this work, we set out to determine whether one can leverage the advantages of CFE to engineer RBCs, both internally and externally. Compared to other methods, CFE is fast, scalable, cheap, and it does not require purification of proteins nor necessitate complex preliminary genetic engineering. We used an all-*E. coli* CFE system, commercially available under the name myTXTL, to engineer mouse RBCs. We show that RBC ghosts can be reloaded with TXTL reactions to execute elementary gene circuits, including a glucose sensor. The cell-free synthesis (CFS) of phages demonstrates that CFE reactions programmed with relatively large DNAs can be achieved inside RBC ghosts. Cell-free synthesized peptides and proteins, fused to a natural RBC-derived transmembrane anchor, can be tightly and durably bound to the outer membrane of undamaged RBCs. We attached a SARS-CoV-2 antigen to the RBC membrane, which was recognized by a specific antibody from a commercial source. Overall, CFE interfaces remarkably well with RBCs and proves effective as an alternative tool for engineering that specific cell type. This hybrid approach bridges CFE, a versatile platform for bottom-up synthetic biology, with RBCs, a physiologically relevant chassis with properties not accessible by other types of synthetic cells, like liposomes.

## Results and Discussion

### 1. CFE and Preparation of RBSCs

We prepared and used two *E. coli* lysate-based CFE systems [84–86], both relying on the endogenous transcription (*E. coli* core RNA polymerase, sigma factor 70) and translation machineries provided by the lysates. The standard CFE system comprises a lysate prepared from the *E. coli* strain BL21-ΔrecBCD Rosetta2, with a recBCD knockout to prevent linear DNA from degrading [87]. The second CFE system comprises a lysate prepared from the *E. coli* strain ClearColi, an endotoxin-free strain derived from BL21 devoid of lipopolysaccharide (LPS), carrying the plasmid Rosetta2. LPS are endotoxins from the outer membrane of Gram-negative bacteria that act as major triggers for inflammation in higher organisms. Except for the lysates, the CFE reactions conducted in this work were the same and based on the TXTL toolbox 2.0 [85]. We used the fluorescent protein mCherry as a quantitative reporter in most experiments, due to its stable fluorescence over a wide physiological pH range [88], its established monomeric state, which unlike many GFP variants-prevent non-specific binding to cellular membranes [89,90], and the low background fluorescence in the red wavelengths compared to other colors. CFE was carried out either from an *E. coli* sigma 70 promoter (P70a-mcherry), from the T7 promoter through a transcriptional activation cascade (P70a-T7rnap → T7-mcherry), or from the P28a promoter through a transcriptional activation cascade (P70a-S28 → P28a-mcherry) as described before [85,86]. In batch mode, mCherry was produced at concentrations of 80-100 µM (2-2.5 mg/ml) from linear DNA (PCR amplicons) or plasmids **(Fig. S1)**. The time course of mCherry synthesis ranged from 10 to 20 hours and was slightly shorter in the ClearColi CFE system. As expected, the standard CFE system lysate contains some endotoxins (lipopolysaccharides, LPS), while the amount of LPS in the ClearColi CFE system lysate was negligible **(Table S1)**.

A CFE reaction is a complex physiological solution containing macromolecules, among which ribosomes and DNA are physically the largest. Ribosomes are globular, with a diameter of about 25 nm, whereas DNA’s size depends on its length. In this study, the smallest dsDNA used for TXTL was on the order of one kbp (P70a-*mcherry*, linear). With a persistence length of about 50 nm (150 bp), a 1-kbp linear dsDNA has a root-mean-square end-to-end distance of 175 nm (worm-like chain) and a radius of gyration of about 75 nm (worm-like chain, effective size 150 nm). Given this, we prepared RBC ghosts by a one-step hypotonic lysis. The RBCs used in this work were isolated from male or female mice following standard procedures and were handled using two buffered solutions, buffer C1 and buffer C2. To prevent RBC lysis, we prepared isotonic buffer solution C1, which included potassium and magnesium necessary for CFE reactions. When observed under the microscope, the RBCs washed and resuspended in buffer C1 had a normal biconcave disk shape **(Fig. S2)**. RBCs had no fluorescence when observed by microscopy in the green channel (eGFP or FITC, Exc. 484 nm, Em. 515 nm) and the red channel (mCherry, Exc. 585 nm, Em. 610 nm) **(Fig. S2)**. RBCs were lysed through a single wash with a hypotonic solution (buffer C2), which contained low concentrations of the salts and biochemical reagents used for CFE reactions. The membranes were isolated by centrifugation and washed extensively with buffer C2 after lysis at 4 °C to prevent resealing. From this point, the RBC membranes were mixed with the CFE reaction to be encapsulated and incubated at 37 °C for 30 minutes to reseal the RBC ghosts and create RBSCs. The RBSCs were washed and resuspended in buffer C1 after encapsulation.

As a preliminary step to visualize the RBC ghosts, we encapsulated a control CFE reaction (lacking DNA) containing 50 µM FITC used as an internal volume marker, while membranes were stained with Nile Red. **(Fig. S3)**. Under the microscope, the RBSCs resuspended in buffer C1 appeared in different shapes, some resembling natural RBCs (biconcave disks), some more elongated or spherical **(Fig. S3)**. Most of the RBCs (> 90%) had both FITC and Red Nile fluorescence. We noticed that the RBSCs reacted to osmotic stress similarly to natural RBCs. Upon a slight increase of the outer solution osmolarity (100 mOsm), more than 75% of the RBSCs spontaneously adopted a spherical shape without lysis or leak of the FITC dye. We used this approach to measure the encapsulation of several dyes because integrating the fluorescence signal of spheres is easier to quantify as a function of the radius, as we reported before using liposomes [91].

We first characterized the encapsulation of two fluorescent probes to confirm that resealed RBSCs can tightly encapsulate molecules. FITC (0.5 kDa, 50 µM) and TRITC-dextran (tetramethyl rhodamine isothiocyanate dextran, 40 kDa, 50 µM) were mixed separately in a CFE reaction without DNA. Each CFE reaction was then encapsulated into RBSCs. The same experiment was done with liposomes, at four different concentrations for each dye (0, 1, 20, 50 µM), which we used to calibrate the encapsulation of each dye. The fluorescence of liposomes was plotted against their radius squared to account for the microscopy geometry and enable linear fits **(Fig. S4, S5)**, as described previously [91]. We used the calibrations to estimate the concentrations of FITC and dextran-TRITC encapsulated in the RBSCs. Two repeats at two different times (1 hour and 24 hours after encapsulation) were carried out **(Fig. S6)**. The results confirmed that neither of the two dyes leaked over a period of at least 24 h. We co-encapsulated both dyes (FITC, 50 µM, and TRITC-dextran, 50 µM) and measured a relatively high degree of fluorescence correlation **(Fig. S6)**.

To characterize the distribution of the RBCs and RBSCs’ sizes, we applied an osmotic stress to convert them to spheres and measured the radii. The RBCs and RBSCs’ radii distribution shows that RBSCs have smaller radii on average **(Fig. S7).** The RBCs’ radii ranged from 2.26 µm to 3.48 µm, while the RBSCs’ radii ranged from 0.65 µm to 1.97 µm. We observed a reduction in the size of the resealed RBC ghosts, in which their diameters appeared to be reduced to approximately half that of native RBCs. It has been previously shown that RBC ghosts used as protein carriers are typically smaller than normal erythrocytes [92], a characteristic known to depend on the experimental settings used during hypotonic lysis and subsequent temperature-induced resealing [93]. Furthermore, it has been demonstrated that the volume of encapsulated solution into RBC ghosts is inversely proportional to the resealing medium osmolarity [94], explaining why the resulting RBSCs are smaller than RBCs when exposed to the CFE reaction’s osmolarity (∼650 mOsm/L), which is larger than the RBC cytosol’s osmolarity (∼300 mOsm/L).

### 2. CFE reactions can be carried out within RBSCs

To measure the concentration of mCherry produced by CFE inside the RBSCs, we first established a calibration of mCherry **(Fig. S8)**, using pure mCherry and following the same procedure as for FITC and TRITC-dextran. Encapsulation of the CFE reactions with DNA was conducted in the same manner as with the dyes **(Fig. 1A)**. Next, mCherry was synthesized inside the RBSCs by four different sets of DNAs: from the *E. coli* promoter P70a, from a T7 promoter through a transcription activation cascade, and from linear amplicons and plasmids. We measured the concentration of mCherry produced after 12 hours of incubation for four DNA concentrations. As observed in batch mode CFE reactions **(Fig. S1)**, mCherry synthesis was slightly greater when plasmids were used **(Fig. 1B, C)**. With an average concentration between 20-40 µM (0.5-1 mg/ml), mCherry synthesis was on the order of two to three times smaller inside the RBSCs than in test tubes, yet strong enough to achieve CFE from DNAs other than reporters. This result agrees with the encapsulation of FITC and TRITC-dextran, whose concentrations inside RBSCs are smaller than in the initial CFE reaction **(Fig. S6)**. Outliers showing either much smaller or much larger mCherry concentrations were present in most of the four sets of experiments. This observation is also supported by the presence of outliers in the case of the encapsulation of dyes **(Fig. S3, S6)**, which show that the encapsulation process can be highly heterogeneous for some RBSCs as it was previously described in liposomes [95,96] or erythrocytes [65,97]. The fluorescence from the synthesized mCherry was easily observed under the microscope **(Fig. 1D)**. The kinetics of mCherry synthesis into the RBSCs lasted about 10 hours **(Fig. 1E)**. The synthesized mCherry using the ClearColi CFE system, via the linear DNA and plasmid versions of the T7 transcriptional activation cascade, produced similar results to the standard CFE system except for a slightly smaller synthesis yield **(Fig. S9)**.

**Figure 1.**
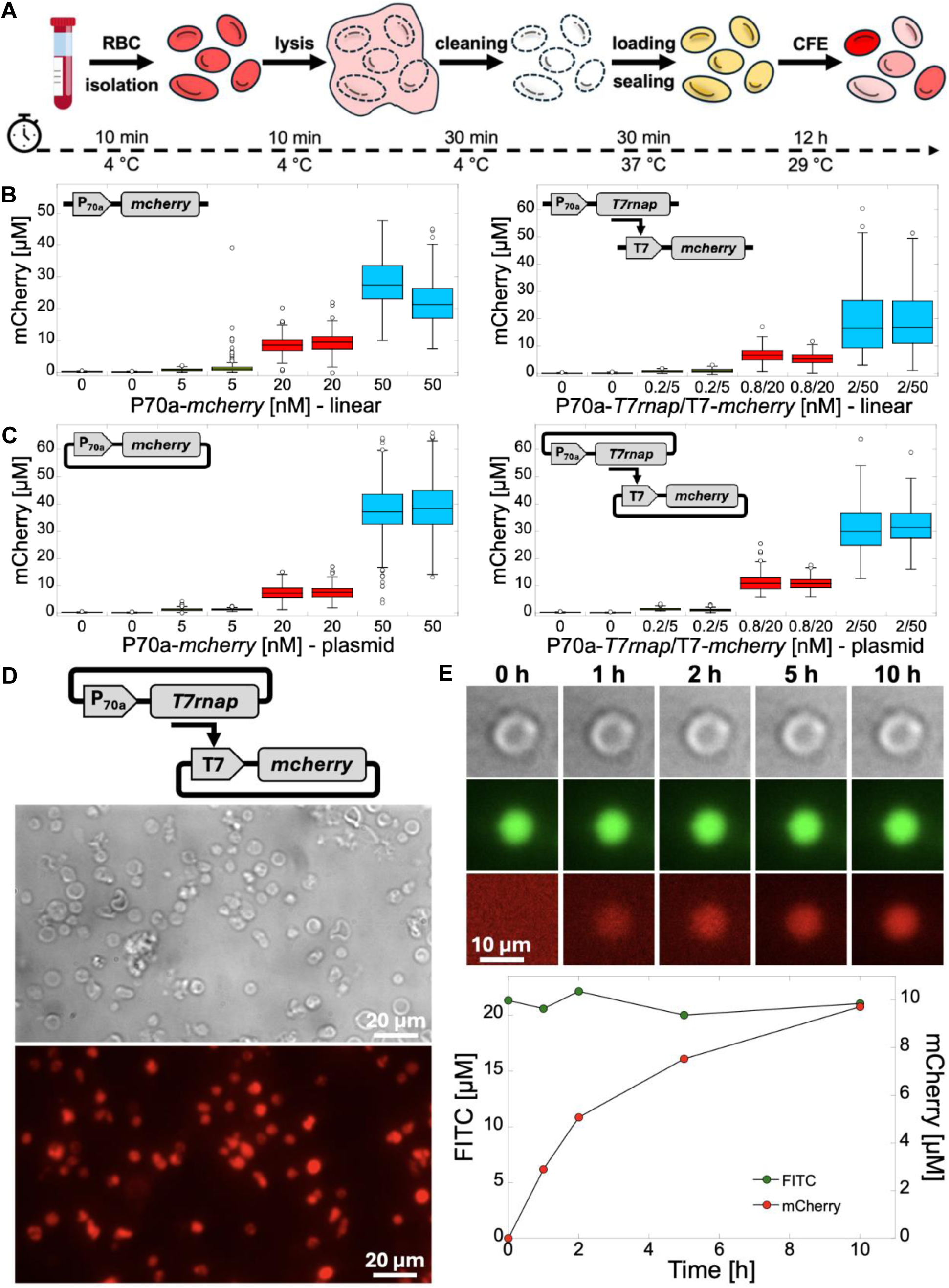
CFE can be achieved inside RBC ghosts. **A.** Diagram of the CFE-based RBC ghosts’ preparation (RBSCs). **B.** Box plots of mCherry cell-free synthesis inside RBSCs using linear DNAs, from a P70a promoter and from a transcriptional activation cascade. Two repeats of each DNA concentration are shown. The fluorescence of two hundred RBSCs was measured and quantified after 12 hours of incubation. **C.** Box plots of mCherry cell-free synthesis inside RBC ghosts using plasmids, from a P70a promoter and from a transcriptional activation cascade. Two repeats of each DNA concentration are shown. The fluorescence of two hundred RBSCs was measured and quantified after 12 hours of incubation. **D.** Fluorescence microscopy images of mCherry cell-free synthesis inside RBSCs. Images acquired after 12 hours of incubation. P70a-*T7rnap* 2 nM, T7-*mcherry* 50 nM. **E.** Kinetics of mCherry cell-free synthesis inside an RBSC supplemented with 20 µM fluorescein (used as a dye). P70a-*T7rnap* 2 nM, T7-*mcherry* 50 nM.

Next, we examined whether the RBSC system could also enable the synthesis of phages. We use the T7 bacteriophage as a model phage because we demonstrated that it can be synthesized by CFE [16,85,98]. It represents a complex set of about sixty genes encoded into a linear dsDNA genome of 40-kbp, much larger than typical gene circuits. The preparation of the RBSCs and encapsulation of the CFE reaction were performed similarly to those with linear DNAs and plasmids for *mcherry* expression. After 12 hours of incubation, the RBSCs were washed, then lysed, and a spotting assay was performed to measure the concentration of T7 phages produced inside the RBSCs. We measured on the order of 10^9^-10^11^ PFU/ml (plaque-forming units per milliliter), demonstrating that RBSCs can accommodate large and complex genetic elements **(Fig. S10)**. No plaques were visible from the supernatant solution containing the RBSCs before lysis. The T7 phage titer was comparable to a batch mode reaction [16,85,98].

### 3. Biosensing can be achieved using RBSCs

Next, we explored whether RBSCs can be functionalized with sensory mechanisms. First, we tested an arabinose-inducible system using an amplification transcriptional activation cascade. The sigma factor 28 gene *S28* was cloned under the arabinose-activating Para promoter [77] into a plasmid bearing the *araC* repressor gene. The *mcherry* gene was cloned under the promoter P28a specific to sigma 28. The two plasmids and the CFE reaction were encapsulated into RBSCs. Induction of mCherry synthesis inside the RBSCs was observed upon addition of 10 mM arabinose to the outer solution **(Fig. 2A)**. To further test the control of arabinose transport into RBSCs, the GLUT1 glucose transporter inhibitor Bay-876 [99], set to 10 µM **(Fig. S11)**, effectively suppressed the induction of mCherry synthesis. This agrees with previous literature, which found that arabinose transport into RBCs, along with several other sugars including glucose, depends on GLUT1 [100]. We found an average RBSC signal ON/OFF ratio (10 mM / 0 mM arabinose signal) of 40 and 44 respectively for the two repeats of this experiment **(Fig. 2B)**.

**Figure 2.**
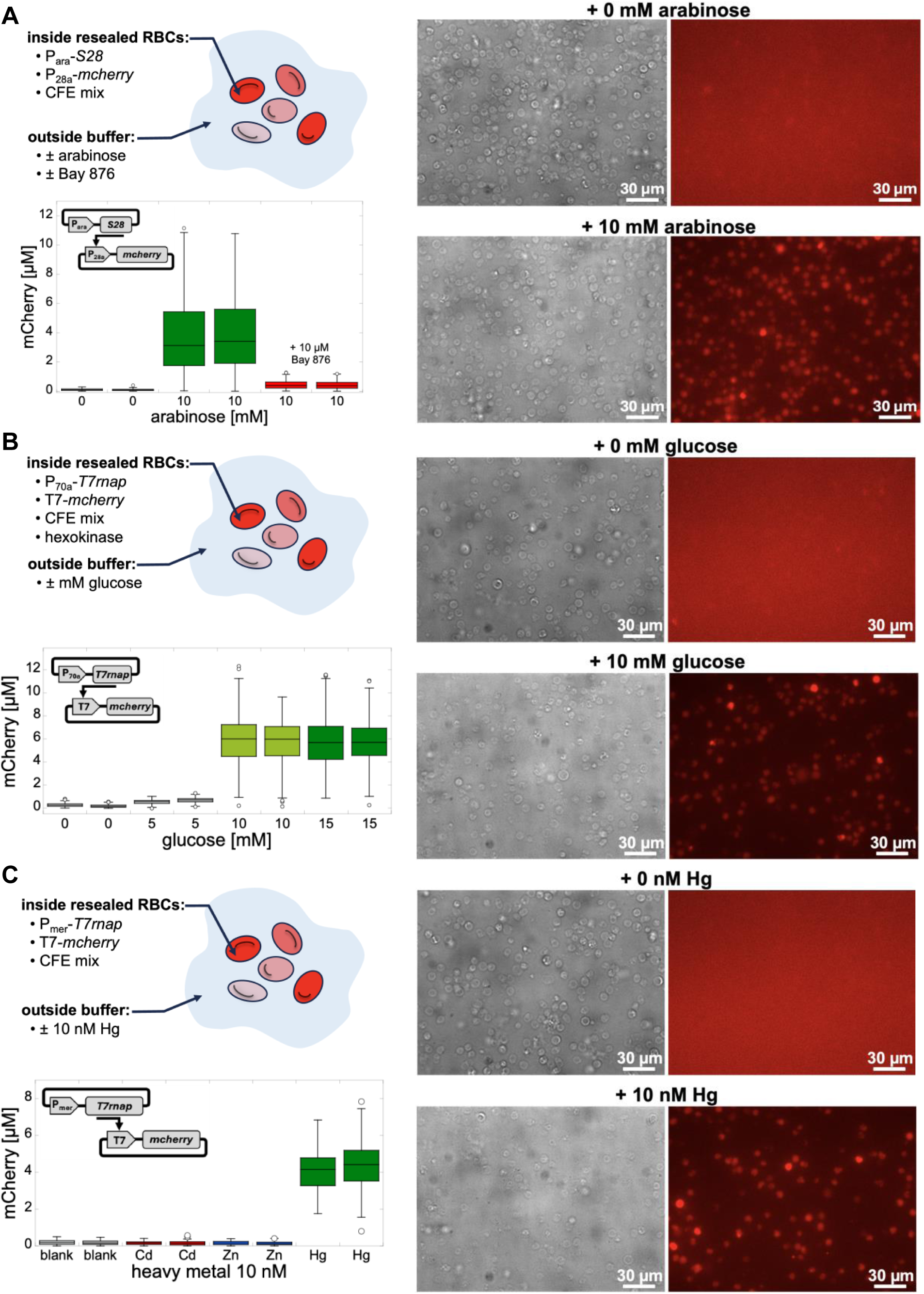
Sensory mechanisms can be carried out using CFE RBSCs. **A.** The arabinose inducible system. Left: diagram of the system using RBSCs. *Sigma factor 28* was expressed through the Para promoter to activate expression of the *mcherry* gene from a P28a promoter. Box plots of mCherry cell-free synthesis inside RBSCs using plasmids. Two repeats of each arabinose concentration are shown. Para-*S28* 2 nM, P28a-*mcherry* 50 nM. The fluorescence of two hundred RBC ghosts was measured and quantified after 12 hours of incubation for each arabinose concentration. Right: Fluorescence microscopy images of mCherry cell-free synthesis inside RBSCs. Images acquired after 12 hours of incubation. Para-*S28* 2 nM, P28a-*mcherry* 50 nM. **B.** A glucose sensor system. Left: diagram of the system using RBSCs. *mcherry* was expressed through the T7 promoter. Box plots of mCherry cell-free synthesis inside RBSCs using plasmids. Two repeats of each glucose concentration are shown. The fluorescence of two hundred RBSCs was measured and quantified after 12 hours of incubation for each glucose concentration. P70a-*T7rnap* 2 nM, T7-*mcherry* 50 nM. Right: Fluorescence microscopy images of mCherry cell-free synthesis inside RBSCs. Images acquired after 12 hours of incubation. P70a-*T7rnap* 2 nM, T7-*mcherry* 50 nM. **C.** A mercury sensor system. Left: diagram of the system using RBSCs. The *T7rnap* gene was expressed through the Pmer promoter to activate expression of the *mcherry* gene from a T7 promoter. Box plots of the cell-free synthesis of mCherry inside RBC ghosts using plasmids. Two repeats of each heavy metal (fixed at 10 nM) are shown. The fluorescence of two hundred RBSCs was measured and quantified after 12 hours of incubation for each heavy metal. Pmer-*T7rnap* 2 nM, T7-*mcherry* 50 nM. Right: Fluorescence microscopy images of mCherry cell-free synthesis inside RBSCs. Images acquired after 12 hours of incubation. Pmer-*T7rnap* 2 nM, T7-*mcherry* 50 nM.

To determine whether the RBSC platform could serve as a biological glucose sensor, we first made a CFE glucose-responsive by adding hexokinase to the reaction. The addition of up to 10 U/ml hexokinase did not affect CFE **(Fig. S12)**. Cell-free synthesis of mCherry as a function of the glucose and hexokinase concentrations was determined in batch mode **(Fig. S12)**. In fasting conditions, a glucose concentration smaller than 100 mg/dL (5.5 mM) is normal, prediabetes occurs at concentrations of 100-125 mg/dL (5.5-7 mM), while diabetes sets in at a glucose concentration greater than 125 mg/dL (7 mM). We prepared RBSCs and measured a response to glucose only at diabetes levels **(Fig. 2B, S13)**, and only in the presence of hexokinase **(Fig. S13)**. mCherry synthesis within RBSCs was clearly visible under the microscope **(Fig. 2B)**. The GLUT1 inhibitor Bay-876 effectively prevented the intake of glucose, as observed by the inhibition of mCherry cell-free synthesis **(Fig. S13)**. We found an average RBSC signal ON/OFF ratio (10 mM / 0 mM glucose signal) of 22 and 34 respectively for the two repeats of this experiment **(Fig. 2B)**.

Lastly, we examined whether RBSCs could also be engineered to sense toxins such as heavy metals. We focused on mercury because a cell-free mercury genetic biosensor has already been demonstrated [101], and RBCs are known to transport heavy metals such as mercury across their membranes [102]. To create an amplification transcriptional activation cascade induced by mercury, we cloned the *T7rnap* gene under the merR-Pmer (plasmid Pmer-*T7rnap*) repressor-promoter operator system. We verified that this system responds specifically to mercury in batch mode CFE reactions **(Fig. S14)**. The concentration of 10 nM heavy metals was chosen for biosensing tests, which corresponds to 2*10^-6^ ppm. In the USA, the FDA has a limit of 1 ppm mercury in foods such as fish and 0.002 ppm in drinking water. A detection limit of 2*10^-6^ ppm is one thousand times smaller than the smallest limit. The RBSC mercury sensor responded as specifically as in batch mode reactions **(Fig. 2C)**. The synthesis of mCherry was clearly visible inside the RBSCs **(Fig. 2C)**, which demonstrated their use as toxin biosensors. We found an average RBSC signal ON/OFF ratio (10 nM / 0 nM mercury signal) of 21 and 25 respectively for the two repeats of this experiment **(Fig. 2C)**.

### 4. Payloads can be attached to the membrane of RBCs using CFE only

Refactoring erythrocytes into genetically programmable synthetic cells enables the creation of CFE-based RBSCs with diverse possible utilizations. Another significant interest for developing biomedical applications is the binding of therapeutically relevant molecular payloads to the RBCs’ outer membrane. Strategies to attach payloads to the membrane of RBCs have been implemented to deliver and target therapeutics in the body, as well as to induce immune tolerance or immunogenicity [62,72,74,75]. RBCs loaded with antigens on their outer membrane can either induce tolerance or an immune response, depending on different factors [103,104], and can also be targeted to particular cells by attaching specific antibodies [105]. First, we used the toxin alpha-hemolysin (AH) to verify that cell-free synthesis in the presence of added washed RBCs to the reaction can be achieved to anchor a payload to the outer membrane of the RBCs. As expected, the fusion protein AH-eGFP colocalized at the RBC membrane **(Fig. S15)**. AH-eGFP induces hemolysis of the RBCs, which take a spherical shape. Based on this result, our goal was to show that one can bind biologically relevant payloads to the outer membrane of RBCs, using CFE only, robustly, and durably without noticeable perturbation to the cell.

First, we searched for a stable, preferably natural, RBC membrane anchor that can be genetically encoded. Glycophorin A (GPA) is the most abundant membrane protein of RBCs. It is bound to the RBC membrane by a single-pass transmembrane hydrophobic alpha-helix. The GPA transmembrane helix was previously synthesized at high yield in *E. coli* [106]. We fused the *mcherry* moiety *sfcherry11* to the GPA’s transmembrane helix (GPAtm) and used a quartz crystal microbalance with dissipation (QCMD) to determine whether cell-free synthesized GPAtm binds to lipid bilayers **(Fig. 3A)**. As demonstrated before, the integration of cell-free synthesized membrane proteins into supported lipid bilayers (SLBs) is characterized by a net drop in the resonance frequency on a QCMD [107]. A net drop in the frequency shift was observed when sfcherry11-GPAtm was dynamically synthesized on a DOPC SLB **(Fig. 3B)**. We repeated this experiment on an *E. coli* lipid bilayer (ECL) and made the same observation **(Fig. S16)**.

**Figure 3.**
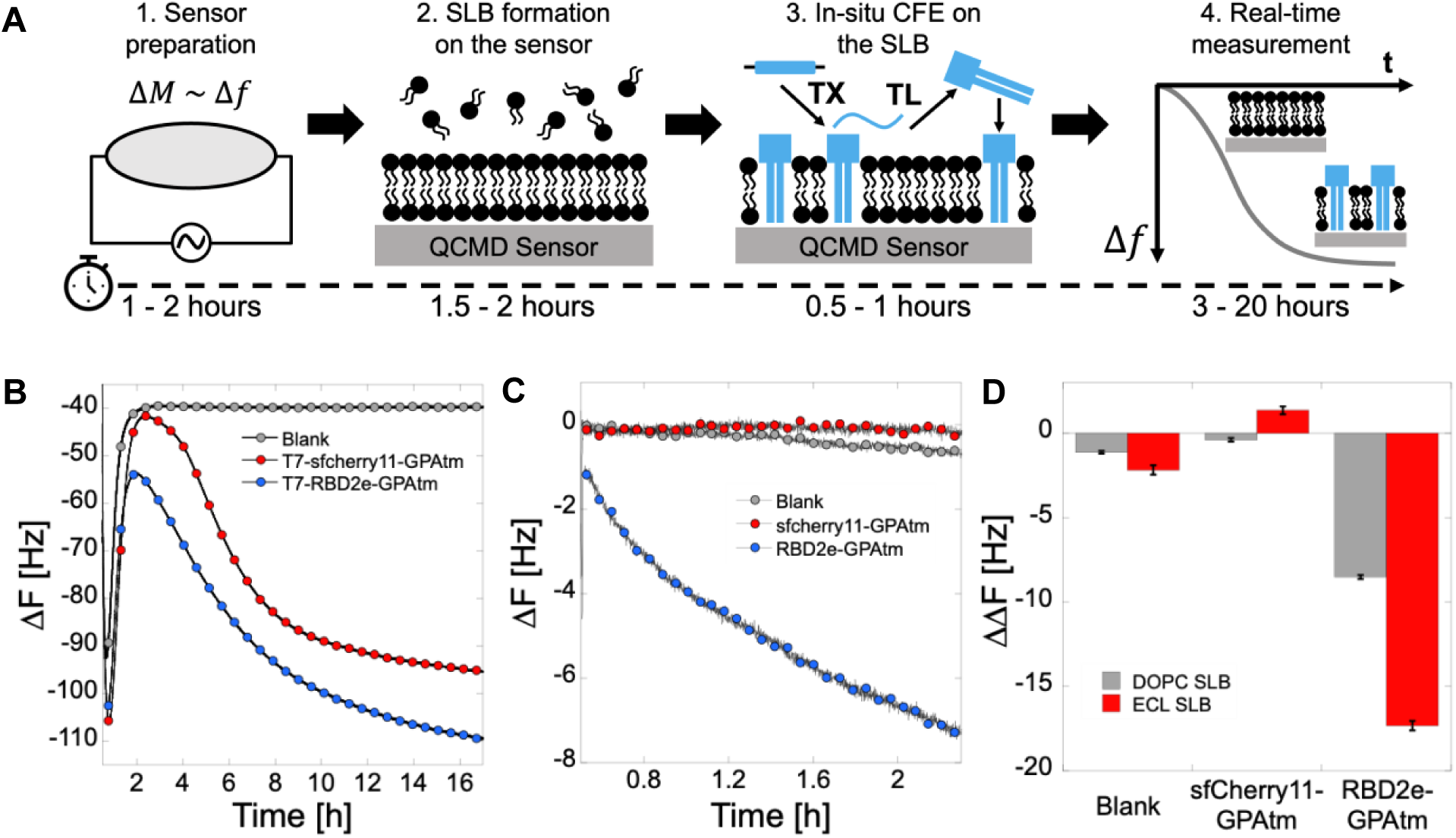
Payloads can be attached to the membrane of RBCs using CFE only. **A.** Diagram of the QCMD workflow. **B.** The transmembrane helix GPAtm enables anchoring payloads to phospholipid bilayers. Cell-free synthesized sfcherry11-GPAtm and RBD2e-GPAtm are both inserted into a DOPC membrane. Blank reaction consisted of a CFE reaction with no added DNA. **C.** The antibody 5G8 (0.5 µM) specific to RBD2e was incubated on three different DOPC supported lipid bilayers (SLBs): a blank SLB, an SLB with sfcherry11-GPAtm and an SLB with RBD2e-GPAtm. **D.** Endpoint of the change in frequency shift for a DOPC and *E. coli* lipid (ECL) SLBs after incubation of the antibody 5G8 (0.5 µM).

Next, we assessed whether we could use GPAtm to bind a viral antigen. Recently, we showed that one can use our *E. coli* CFE system to synthesize the receptor binding domain of the S1 spike protein of SARS-CoV-2 (RBD2) [108]. Specifically, we focused on segment e (RBD2e), which corresponds to amino acids 321-420 of the S1 spike protein. This epitope is recognized by the antibody 5G8, which is commercially available. We made an *RBD2e-GPAtm* construct and observed its integration in both DOPC and ECL membranes **(Fig. 3B, S16)**. Then, specific binding of the antibody 5G8 was verified on the QCMD. First, we integrated sfcherry11 - GPAtm or RBD2e-GPAtm separately into either DOPC or ECL SLBs. The QCMD modules were washed with buffer, and 0.5 µM of the antibody 5G8 was incubated on the SLBs. A net drop in frequency shift was observed only for SLBs with RBD2e-GPAtm **(Fig. 3C, 3D, S16)**. To confirm this result, we integrated RBD2e-GPAtm into the membrane of RBCs by CFE then washed the RBCs **(Fig. 4A)** and noticed no significant morphological changes to the cells. The RBCs were incubated with 0.5 µM of the antibody 5G8, followed by extensive washing. On an SDS-PAGE, we observed the presence of the antibody 5G8 only when it was incubated with the RBD2e-GPAtm RBCs **(Fig. 4B)**. We repeated the same experiment with a ClearColi CFE system and obtained the same results **(Fig. 4C).**

**Figure 4.**
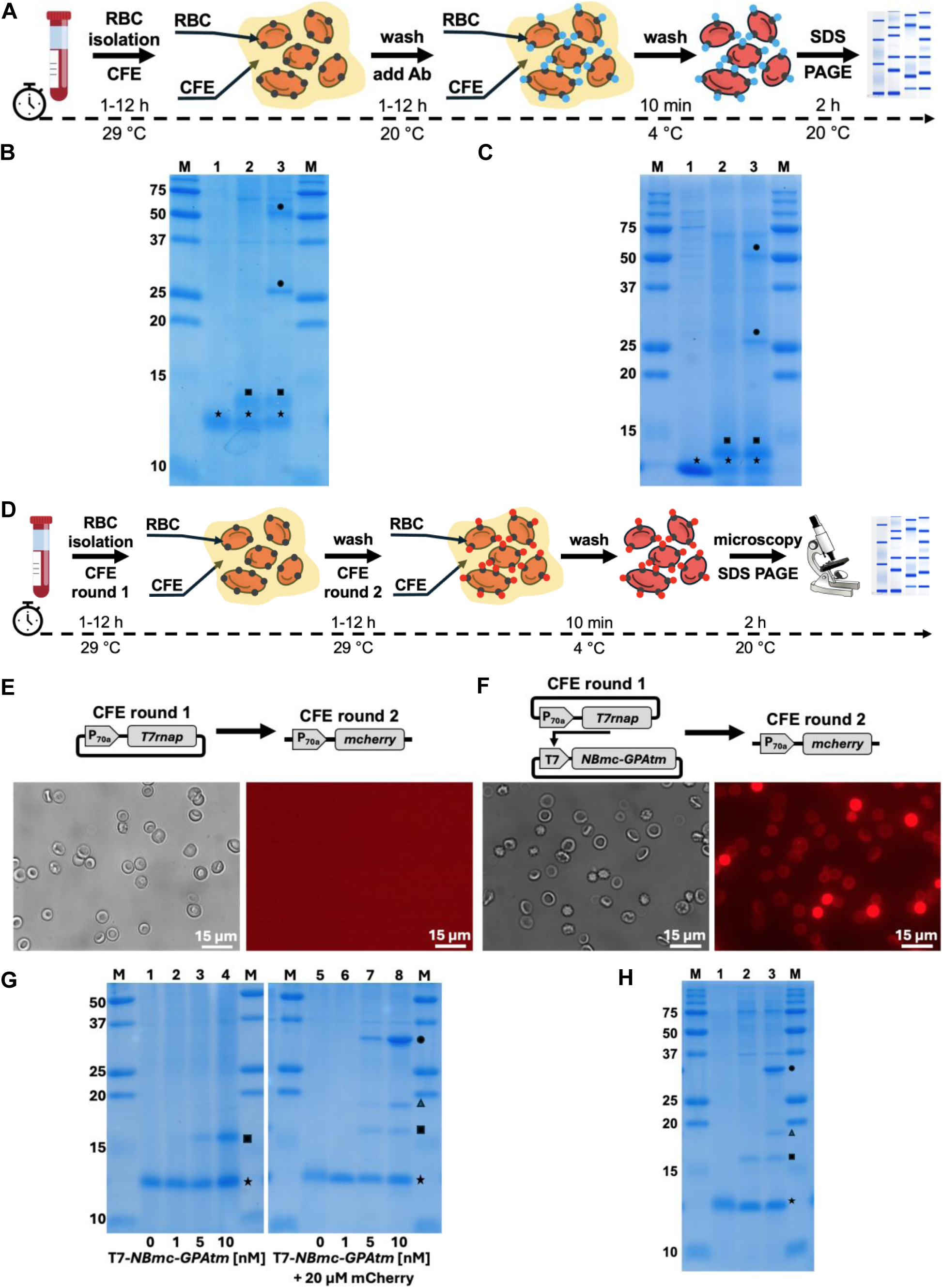
Payloads can be attached to the membrane of RBCs using CFE only. **A.** Diagram of the assay of the RBD2e antigen binding to the outer membrane of RBCs and its subsequent interaction with the 5G8 antibody. **B.** SDS PAGE of RBCs. Symbols: star (hemoglobin, 14 kDa), square (RBD2e-GPAtm, 15 kDa), circle (5G8, large and small subunits). M: marker, molecular mass in kDa on the left of the gel. 1: RBCs only, 2: RBC + RBD2e-GPAtm, 3: RBC + RBD2e-GPAtm + 5G8. **C.** Same as in B but with a ClearColi CFE system. **D.** Diagram of the assay of mCherry nanobody (NBmc-GPAtm) binding to the outer membrane of RBCs and its subsequent interaction with mCherry. **E.** Microscopy images (phase contrast and fluorescence) of the binding of mCherry when the nanobody NBmc-GPAtm is not synthesized. **F.** Microscopy images (phase contrast and fluorescence) of the binding of mCherry when the nanobody NBmc-GPAtm is synthesized. **G.** SDS PAGEs of the mCherry nanobody interaction onto the outer membrane of RBCs. SDS PAGE on the left: symbols: star (hemoglobin, 14 kDa), square (NBmc-GPAtm, 22 kDa). M: marker, molecular mass in kDa on the left of the gel. 1: RBCs only, 2, 3, 4: RBC + NBmc-GPAtm (three concentrations of T7-*NBmc-GPAtm*). SDS PAGE on the right (20 µM of pure mCherry was added after binding of NBmc-GPAtm to RBCs): symbols: star (hemoglobin, 14 kDa), square (NBmc-GPAtm, 22 kDa). Triangle and circle: mCherry. M: marker, molecular mass in kDa on the left of the gel. 1: RBCs only, 2, 3, 4: RBC + NBmc-GPAtm (three concentrations of T7-*NBmc-GPAtm*). **H.** SDS PAGE of the mCherry nanobody interaction onto the outer membrane of RBCs using a ClearColi CFE system. Symbols: star (hemoglobin, 14 kDa), square (NBmc-GPAtm, 22 kDa). Triangle and circle: mCherry. M: marker, molecular mass in kDa on the left of the gel. 1: RBCs only, 2: RBC + NBmc-GPAtm (one concentration of T7-*NBmc-GPAtm*), no mCherry added, 3: RBC + NBmc-GPAtm (one concentration of T7-*NBmc-GPAtm*), when 20 µM pure mCherry is added.

To show that one can achieve an inverted biomolecular interaction scheme, we attached a nanobody specific to mCherry (NBmc) onto the RBCs. The *NBmc-GPAtm* was expressed in the presence of RBCs. Following a washing step, the RBCs were incubated in a CFE reaction expressing *mcherry* **(Fig. 4D)**. The RBCs were subsequently washed and observed under the microscope. A net fluorescence at the surface of the RBCs was observed, again without noticeable cell morphology changes from the CFE processing **(Fig. 4E, 4F)**. To confirm the specificity of this interaction and show that one can tune the amount of attached antigen onto the RBCs, we repeated this experiment by expressing *NBmc-GPAtm* at four different DNA concentrations (0, 1, 5, 10 nM linear T7-*NBmc-GPAtm*, 2 nM P70a-*T7rnap* plasmid). On an SDS PAGE, one can see different amounts of NBmc-GPAtm attached to the RBCs, and similarly different amounts of mCherry bound to the RBCs **(Fig. 4G)**. No binding occurred at 0 and 1 nM T7-*NBmc-GPAtm*. Note that on the SDS PAGE, several bands appeared for mCherry, which has been described previously [109,110]. The same experiments (RBD2e-GAPtm + 5G8, NBmc + mCherry) were repeated with similar results using the ClearColi CFE system **(Fig. 4H)**. No LPS were detected in these experiments **(Table S2)**. The use of CFE to durably attach antigen payloads to the outer membrane of RBCs is a rapid design-built-test cycle, taking approximately 2 – 25 hours, with no perturbation to RBC morphology.

Although the theoretical molecular weight of the NBmc-GPAtm nanobody is 22 kDa, its band size appeared to be approximately 16–17 kDa on SDS-PAGE. This anomalous migration is typical for stable single-domain nanobodies and hydrophobic membrane fusions. First, the highly stable nanobody core can maintain a compact conformation that migrates relatively further through the gel if there was incomplete denaturation [111]. Second, it has been shown that transmembrane domains, such as GPAtm, bind excessive amounts of SDS and alter protein migration in gels, which can result in a smaller apparent molecular weight [112–114].

### 5. Discussion and Perspectives

This work shows that an *E. coli* CFE system interfaces remarkably well with RBCs. CFE can be used to transform RBCs into genetically programmable synthetic cells that are responsive to their environment, and the RBCs’ outer membrane can be engineered to carry therapeutically relevant payloads. The CFE approach of engineering the cytoplasm of RBCs is novel, and the ability to execute DNAs via CFE inside RBC ghosts shows that nothing, including proteases and nucleases, prevents transcription-translation from occurring in resealed erythrocytes. The engineering of the outer membrane of RBCs using CFE is complementary to previously reported approaches and offers several advantages. It can be achieved using a natural, strong, and permanent membrane anchor (GPAtm). The amount of anchored payload can be tuned. It is fast, cheap, and free of the immunogenic endotoxin LPS. We anticipate that engineering the outer membrane of RBCs without LPS will facilitate *in vivo* applications, avoiding unintended immune responses from CFE modification. Importantly, this approach does not require pre-genetic modifications of the cells. In fact, the CFE engineering of the RBC outer membrane could be used to test whether genetically programmable payloads can be synthesized in an active form and attached to the outer membrane of RBCs before deciding on more labor-intensive genetic modification of erythrocytes. Lastly, the CFE approach to RBC membrane engineering somewhat resembles the Kode technology employed to create kodecytes, except that the lipid moiety is replaced by GPAtm.

Whether other types of cells than RBCs can be engineered using CFE remains to be tested. RBCs are well predisposed to CFE engineering due to their biological and physical properties, their lack of a nucleus, and the ease of preparing ghosts. Because of their similarity, cells modified by the Kode technology could be the first tested with the CFE approach. The Kode technology has been used to modify a wide variety of cells, including white blood cells, platelets, embryos, spermatozoa, epithelial cells, and RBCs [115–118] for applications as diverse as labeling [117], attaching infectious markers [119], modifying cell adhesion, interaction, or changing cell separation and immobilization [115]. It would be interesting to determine whether GPAtm could replace the lipid moiety as a universal membrane anchor, thereby enabling a polypeptide payload attachment method.

Determining the immune system tolerance and response, in mice, for instance, of the RBSCs described in this work is among the next steps to achieve. Do the CFE-based LPS-free RBSCs escape the immune system, or are inflammatory markers elicited? Can we use this system for antigen-specific immunological tolerance? Can we use this system to induce antibody responses? These questions will be addressed in future work.

## Materials and Methods

### Cell-free transcription-translation

Two CFE systems were prepared and used for this work: one from the *E. coli* strain BL21-ΔrecBCD Rosetta2, with a recBCD knockout to prevent the degradation of linear DNA [87], and one from the *E. coli* strain ClearColi (Research Corporation Technologies), an endotoxin-free strain derived from BL21. The plasmid Rosetta2 was transformed into ClearColi before lysate preparation. Except for the lysates, the CFE reactions carried out in this work were the same and based on the TXTL toolbox 2.0 [85]. Cells were grown in 2xYT medium supplemented with phosphates, pelleted, washed, and lysed using a pressure cell press. The lysates were centrifuged, and the supernatant was incubated at 37 °C for 80 minutes. After a second centrifugation, the supernatants were dialyzed at 4 °C for 3 hours. Following a final centrifugation step, the lysates were aliquoted and stored at -80 °C. The volume of the CFE reactions consisted of 33% cell extract and 67% of the other components (energy mix and amino acids). The reaction buffer included 50 mM Hepes (pH 8), 1.5 mM ATP and GTP, 0.9 mM CTP and UTP, 0.26 mM coenzyme A, 0.33 mM NAD, 0.75 mM cAMP, 0.068 mM folinic acid, 1 mM spermidine, 30 mM 3-phosphoglyceric acid (3-PGA), 1 mM dithiothreitol (DTT), 1.5% PEG8000, and 20-40 mM maltodextrin. Amino acids were added at concentrations between 1.5 mM and 6 mM for each of the 20 amino acids. Magnesium (2–5 mM) and potassium (50-100 mM) concentrations were calibrated for mCherry synthesis and T7 phage production to obtain reproducible batches. Reactions were incubated at either 15 °C (cell-free synthesis of RBD2e-GPAtm in batch mode CFE reactions), 22 °C (all QCMD experiments), or 30 °C (default temperature), depending on the application, in either 1.5 mL tubes, 96-well plates, QCMD wells, or encapsulated within RBCs. Fluorescence data from CFE reactions (Agilent Biotek H1 or Neo2) were processed with Kaleidagraph. Other reagents used in this work: hexokinase (Sigma H6302), Bay-876 (VWR T3713), arabinose (Sigma A3256), glucose (Sigma G8270), mercury acetate (Sigma 83352), zinc chloride (Sigma 229997), cadmium acetate (Sigma 229490).

### Blood collection and RBC isolation

Whole blood was obtained from 8 to 12-week-old mice. To prevent coagulation during storage and subsequent handling, 10 mM EDTA was added to the collected blood. Samples were stored at 4 °C and processed within two weeks after collection. RBCs were isolated and washed according to the following RBC preparation procedure.

### RBC ghosts’ preparation for internal CFE

RBC ghosts were prepared and resealed as follows. 10 µL of RBCs were washed in 1 mL of isotonic buffer C1 (Tris-HCl 45 mM, K-Glu 270 mM, Mg-Glu 25.5 mM at pH 8) and collected by centrifugation at 500–1000 × g for 5 minutes. For each experimental condition, 3.5 µL of the resulting RBC pellet was lysed by resuspension in 1 mL of ice-cold hypotonic buffer C2 (K-Glu 1 mM, Mg-Glu 1 mM, CaCl2 1 mM, amino acids 0.02 mM each, Maltodextrin 0.15 mM, PEG8000 0.015%, 3PGA 0.2 mM, NAD 0.001 mM, ATP 0.01 mM, GTP 0.01 mM, UTP 0.01 mM, CTP 0.01 mM at pH 8) and incubated on ice for 10 minutes. The resulting RBC membranes (ghosts) were isolated by centrifugation at 10,000 × g for 10 minutes at 4 °C. The supernatant was discarded, and the pellet was washed once with the same hypotonic buffer under the same centrifugal conditions. To initiate the resealing process, the ghost membranes were resuspended in 25 µL CFE reaction for each condition. The suspension was incubated in a water bath at 37 °C for 30 minutes to allow the membranes to reform intact vesicles. Following incubation, the resealed ghosts were pelleted at 10,000 × g for 10 minutes at 4 °C. The pellet was washed with 1 mL of buffer C1 to remove debris and unsealed membranes, then centrifuged again. Finally, the resealed ghosts were resuspended in 10 µL of buffer C1 and incubated at 29 °C overnight for CFE.

### RBC preparation for membrane engineering

Mature RBCs isolated from blood were prepared as follows. 10 µL of whole mouse blood was washed in 1 mL of buffer C1 and collected by centrifugation at 500–1000 × g for 5 minutes. This washing step was performed twice to ensure complete removal of plasma components, with the supernatant discarded after each cycle. Following the final wash, 5 µL of the resulting RBC pellet was combined with 20 µL of a CFE reaction mixture with the desired DNA. The suspension was then incubated overnight at 29 °C to achieve membrane functionalization. The next day, the RBCs were washed with buffer C1 several times and resuspended in buffer C1 before being observed under the microscope.

### COVID-19 antibody membrane binding assay

Binding of the COVID-19 antigen RBD2e to the RBCs’ membrane was performed by expressing the antigen RBD2e [108] fused to the transmembrane segment of Glycophorin A (P70a-*T7rnap* 0.2 nM, T7p14-*RBD2e-GPAtm* 5 nM, overnight incubation at 29 °C). Following gene expression, the functionalized RBCs were subjected to 4–5 washing cycles with buffer C1 at 4 °C. For each wash, the cells were recovered by centrifugation at 5,000 × g for 5 minutes. For immunodetection, the washed RBCs were incubated overnight at 4 °C with a commercial anti-SARS-CoV-2 5G8 antibody (Abcam, ab-277628), diluted 1:10 in buffer C1. The following day, the RBCs were washed several times in buffer C1 (5,000 × g for 5 minutes) to remove unbound antibodies. The washed RBCs were resuspended in 5 µL of molecular-grade water to induce osmotic lysis. The RBC membranes were recovered by centrifugation (5,000 × g for 5 minutes). The samples were mixed with an SDS-PAGE loading buffer for SDS-PAGE electrophoresis.

### mCherry membrane binding assay

Binding of the mCherry reporter protein to the RBCs’ membrane was performed by expressing the mCherry-specific nanobody *NBmc* [120] fused to the transmembrane segment of Glycophorin A (P70a-*T7rnap* 0.2 nM, T7p14-*C11aCherry-GPAtm* 5 nM, overnight incubation at 29 °C). Following gene expression, the functionalized RBCs were subjected to 4–5 washing cycles with buffer C1 at 4 °C. For each wash, the cells were recovered by centrifugation at 5,000 × g for 5 minutes. For attachment of mCherry to the membrane, the washed RBCs were resuspended in a CFE reaction expressing mCherry (P70a-*T7rnap* 0.2 nM, T7p14-*mCherry* 5 nM, 3 hours incubation at 29 °C). Following mCherry expression, the RBCs were washed several times in buffer C1 (5,000 × g for 5 minutes) to remove unbound mCherry and resuspended in buffer C1 before being observed under the microscope. For SDS-PAGE electrophoresis, some of the samples were mixed with an SDS-PAGE loading buffer.

### SDS PAGE

The protein profile and membrane integration were analyzed by sodium dodecyl sulfate-polyacrylamide gel electrophoresis (SDS-PAGE) using a 15% resolving gel. Lysed RBC samples were mixed with a 2x Laemmli loading buffer (containing 10% SDS and 5% Beta-mercaptoethanol) and denatured at 95 °C for 10 minutes. The 15% resolving gel was prepared with acrylamide/bis-acrylamide (15% v/v, 30%:0.8% ratio), Tris-HCl (375 mM, pH 8.8), SDS (0.1% w/v), ammonium persulfate (0.1% w/v), and TEMED (0.04% v/v). The 4% stacking gel consisted of acrylamide/bis-acrylamide (4% v/v), Tris-HCl (125 mM, pH 6.8), SDS (0.1% w/v), ammonium persulfate (0.1% w/v), and TEMED (0.1% v/v). Electrophoresis was performed in a 1x running buffer (25 mM Tris-base, 192 mM glycine, and 0.1% w/v SDS, pH 8.3) at a constant voltage of 120 V until completion. Following electrophoresis, the gel was rinsed three times with deionized water. The gel was then incubated in Coomassie Brilliant Blue staining solution (Coomassie Brilliant Blue R-250 (0.1% w/v), methanol (40% v/v), and glacial acetic acid (10% v/v)) for 60 minutes under gentle agitation. The staining solution was replaced with deionized water, and the gel was washed under gentle agitation.

### Phage spotting assay

Cell-free T7 phage production (genome from Boca Scientific, 310025) within RBSCs was quantified using the phage spotting assay. Individual RBSC radii were measured under the microscope, and assuming spherical geometry, the volume of each RBC ghost was calculated. The total encapsulated CFE reaction volume was then calculated by summing the volumes of all RBSC in the well. PFU counts were divided by this total volume, yielding PFU per nanoliter. For the phage spotting assay, 1.5% agar-LB plates were pre-incubated at 37 °C for 1 hour. 10 mL of 0.7% soft agar was kept at 55 °C in a water bath. 100 µL of an overnight bacterial culture was mixed with the soft agar and vortexed gently. The soft agar was slowly dispensed onto the LB agar plates to cover the entire surface of the agar plate uniformly. The soft-agar plates were left at room temperature for 15 minutes on a flat surface to solidify. Serial ten-fold dilutions in LB of either cell-free phage reaction or clarified phage lysates were prepared in 200 µL. Spotting: for each phage dilution, 3.5 µL were dropped onto the soft agar. For negative control CFE reactions, the whole reaction was diluted in LB at a final volume of 25 µL and spotted in one droplet onto the soft agar layer. After spotting, the plate was left for 15 minutes on the bench to let droplets absorb onto the soft agar. The plates were incubated at 37 °C, facing down, for 4 hours. Plaques were counted at the dilution where the plaque number was smaller than 20 per spot. Titers were calculated from three serial dilution spots. Plaque uncertainties are estimated at the time of counting (duplication or plaque belonging to the same spot).

### Microscopy

Microscopy images were acquired using an Olympus IX81 inverted microscope controlled by Metamorph software using the proper set of filters for green fluorescence (FITC, eGFP, exposure time: 500 ms), and red fluorescence (TRITC and mCherry, exposure time: 500 ms), bright field (exposure time 20 ms), and phase contrast (exposure time 20 ms). RBCs were imaged in 384-well plates, mounted on a custom-made microplate stage. Time-lapse acquisitions across multiple wells were performed simultaneously. The temperature was maintained at 30 °C by adjusting the temperature of the microscope room. Image processing, including segmentation and particle analysis, was performed with ImageJ. Exported data were plotted using Python scripts.

### Calibration of fluorescence intensity

Calibration of fluorescence intensity was performed to quantify the concentrations of FITC (Sigma F3651), TRITC-dextran 40 kDa (Sigma 42874), and mCherry (Abcam, ab199750), encapsulated or synthesized (mCherry) in the resealed RBC ghosts, using a method described previously [85,86]. The calibration was carried out between 0 µM and 50 µM and found to be linear in this range. First, each of the three dyes was encapsulated at four different concentrations (0, 1, 20, 50 µM) in cell-sized liposomes **(Fig. S4, S5, S8)**. The intensity of about a hundred liposomes was plotted as a function of the square of the radius for each liposome (40x objective) to take into consideration the geometry of the fluorescence acquisition and get a linear fit. We used the calibrations to measure the concentration of FITC, TRITC-dextran (40 kDa), and mCherry inside the resealed RBCs. For this, we applied a slight osmotic pressure (hypertonic, 10-20 mOsm) onto the RBCs to change their shape to spheres and thus use the calibrations obtained from spherical liposomes **(Fig. S3)**. This osmotic stress did not destabilize the RBCs.

### Emulsion transfer method for liposomes preparation

All the phospholipids were obtained from Avanti Polar Lipids in powder form. The liposomes were prepared using a lipid composition of POPC (850457P, 760.08 g/mol) and PEG-PE 5000 (880200P, 5745.03 g/mol). In this work, the composition used was POPC/PEG-PE 80/20 (molar ratio) at a total lipid concentration of 100 µM. This corresponds to 80 µM of POPC (42 µg/mL) and 20 µM PEG-PE (85 µg/mL). Lipids (POPC, PEG-PE) were dissolved in chloroform at appropriate concentrations (100 mg/mL for POPC and 10 mg/mL for PEG-PE) and stored in airtight glass vials with Teflon-lined caps at −20 °C. Lipid stocks are stable for up to one month at −20 °C in the dark. In a 7 mL glass vial, lipids are mixed in 200 μL hexadecane. Typically, 5 μL of POPC and 5 μL of PEG-PE were added and vortexed for 15 seconds. 1 mL of long-chain mineral oil (Wako liquid paraffin) was added, and the mixture was heated for 1–2 hours at 70 °C, vortexed briefly to ensure homogeneity, and used directly. In a microcentrifuge tube, 200 μL of the lipid-in-oil mixture was combined with 10 μL of the CFE reaction containing the desired components. The solution was emulsified by vortexing or rubbing on a rack, producing CFE droplets encapsulated in lipid micelles. The emulsion was transferred onto a new tube containing 250 μL of buffer C1 and centrifuged at 5000 x g for 10 minutes to form liposomes. After centrifugation, the oil phase is removed. 10 µL of liposome pellet is transferred to a new 1.5 ml tube containing 90 µL of buffer C1. In a 384-well plate, 90 μL of buffer C1 was combined with 10 μL of liposomes for further observation.

### QCMD measurements

A Nanoscience QSense Analyzer (Biolin Scientific, Gothenburg, Sweden, four-channel quartz crystal microbalance with dissipation monitoring, open modules) was used to verify the binding of RBD2e-GPAtm to supported lipid bilayers (SLBs) formed from DOPC (1,2-dioleoyl-sn-glycero-3-phosphocholine, Avanti Lipids, 850375P) or ECL (*E. coli* lipids, Avanti Lipids, 100500P) and to confirm the specificity of binding of the commercial anti-SARS-CoV-2 5G8 antibody (Abcam, ab-277628) onto the RBD2e antigen anchored to SLBs (RBD2e-GPAtm) adsorbed on the sensors (QSense QSX303 SiO2 sensors). The QCMD experiments were conducted as previously described [107]. The SLBs were formed on the sensors using the Solvent Assisted Lipid Bilayer (SALB) method. The sensors were cleaned and locked into the modules. 800 µL of Tris NaCl buffer (10 mM Tris, 150 mM NaCl, pH 7.5) was added to the open module wells onto the sensors and incubated at RT for 10 minutes. To avoid sensor dewetting, transitions between buffers or solutions during SALB were sequentially replaced using the following Standard Exchange protocol (SE) that consisted of four repeats of the following cycle: (i) volume of the solution in each well was removed such that 130 µL remained, (ii) 800 µL of the new solution was added, and (iii) mixed via pipetting. For the SALB final step of the transition to isopropanol, 400 µL isopropanol was added to the 130 µL remaining liquid, then left to incubate at 22 °C for 5 minutes. Lipids used for SLBs (kept at -20 °C in isopropanol) were briefly heated at 55 °C on a dry heat block, gently vortexed, mixed with isopropanol, and added to the wells on the sensors at the following final concentrations: 3 mg/mL (ECL) and 1.5 mg/mL (DOPC). Once added, lipid isopropanol mixtures were left to incubate on the sensors at 22 °C for 25 minutes for ECL and 60 minutes for DOPC. After lipid isopropanol incubation, the lipid solution in each well was removed, and four SE cycles were completed using Tris NaCl buffer. Each well was left with 930 µL Tris NaCl and incubated at 22 °C for 15 minutes to stabilize the SLBs. Before the addition of the CFE reaction, all liquid was removed. 100 µL of CFE reaction was added directly to the sensor surface. The open module was taped closed with transparent adhesive tape (non-permeable to water vapor) and left to incubate at 22 °C for at least 16 hours. After the CFE reaction, the QCMD wells were cleaned using four SE cycles with Tris NaCl buffer at 22 °C and left to incubate for 15 minutes, followed by a cleaning step with buffer C1 (22 °C for 20 minutes). All liquid was removed, then 100 µL of 10 µg/mL anti-SARS-CoV-2 5G8 antibody in buffer C1 was added to the sensors and incubated for 4 hours. The wells were flushed with buffer C1, incubated for 20 minutes at 22 °C, then washed with Tris NaCl buffer (10 mM Tris, 150 mM NaCl, pH 7.5, 22 °C for 15 minutes).

### LPS measurements

The concentration of Lipopolysaccharides (LPS) within the cell-free reactions was quantified using the ToxinSensor™ Chromogenic LAL Endotoxin Assay Kit (GenScript), following the manufacturer’s instructions. To evaluate the presence of LPS on RBC membranes, cell-free reactions expressing mCherry were performed in the presence of RBCs, as described above. Following a 24-hour incubation, the RBCs were recovered and subjected to ten successive wash cycles in buffer C1 to remove any non-specifically bound components. After the final wash, the RBC pellet was resuspended in LAL reagent water (provided with the kit) to release membrane-associated LPS for subsequent chromogenic LAL analysis.

### DNA

All the DNA sequences are reported in **Supplementary File XX**. We either used plasmids or linear DNAs. DNAs were obtained either by PCR or purchased (Twist Biosciences, IDT). DNA stock solutions were quantified with a spectrophotometer (ThermoFisher Scientific, NanoDrop 2000). Genes were expressed either from the strong *E. coli* promoters P70a, or from the P28a or the T7 transcriptional activation cascades.

## Data Availability

The data underlying this article will be shared on reasonable request to the corresponding author.

## Supplementary Data

Supplementary Information file: 16 figures, 2 tables Supplementary Data file: DNA sequences

## Acknowledgements

We thank the University of Minnesota Research Animal Resources for providing the mouse blood samples. We thank Frank Perez from the Institut Curie (Paris, France) for providing us with the DNA encoding for the mCherry nanobody. We thank Aset Khakimzhan for technical assistance.

## Author Contributions

David Garenne and Seth Thompson performed the experiments and edited the manuscript. Vincent Noireaux supervised the work, wrote the manuscript, and acquired funding.

