## Supplementary material for "Engineering erythrocytes into synthetic cells using cell-free gene expression": SI

David Garenne, Seth Thompson, Vincent Noireaux
School of Physics and Astronomy, University of Minnesota, Minneapolis, MN 55455, USA


| Item | Title | Pages |
|---|---|---|
| Figure S1 | CFE in batch mode | 2 |
| Figure S2 | Microscopy images of washed mouse RBCs | 3 |
| Figure S3 | Preparation and labeling of RBSCs | 4 |
| Figure S4 | Calibration of fluorescein dye (FITC) intensity into RBSCs | 5 |
| Figure S5 | Calibration of dextran-TRITC dye intensity into RBSCs | 6 |
| Figure S6 | Quantification of dyes encapsulation into RBSCs | 7 |
| Figure S7 | Probability distribution of the RBCs and RBSCs radii | 8 |
| Figure S8 | Calibration of mCherry intensity into RBSCs | 9 |
| Figure S9 | CFE of *mcherry* into RBSCs using e ClearColi CFE system | 10 |
| Figure S10 | Cell-free synthesis of phage T7 inside RBSCs | 11 |
| Figure S11 | Effect of Bay-876 on CFE | 12 |
| Figure S12 | Effect of hexokinase on CFE | 13 |
| Figure S13 | A glucose sensor system | 14 |
| Figure S14 | CFE using the Pmer system in batch mode CFE reactions | 15 |
| Figure S15 | CFE of alpha-hemolysin (AH) in the presence of RBCs | 16 |
| Figure S16 | Anchoring payloads to phospholipid bilayers using GPAtm (QCMD) | 17 |
| Table S1 | Concentration of LPS in lysates | 18 |
| Table S2 | Concentration of LPS after CFE | 18 |

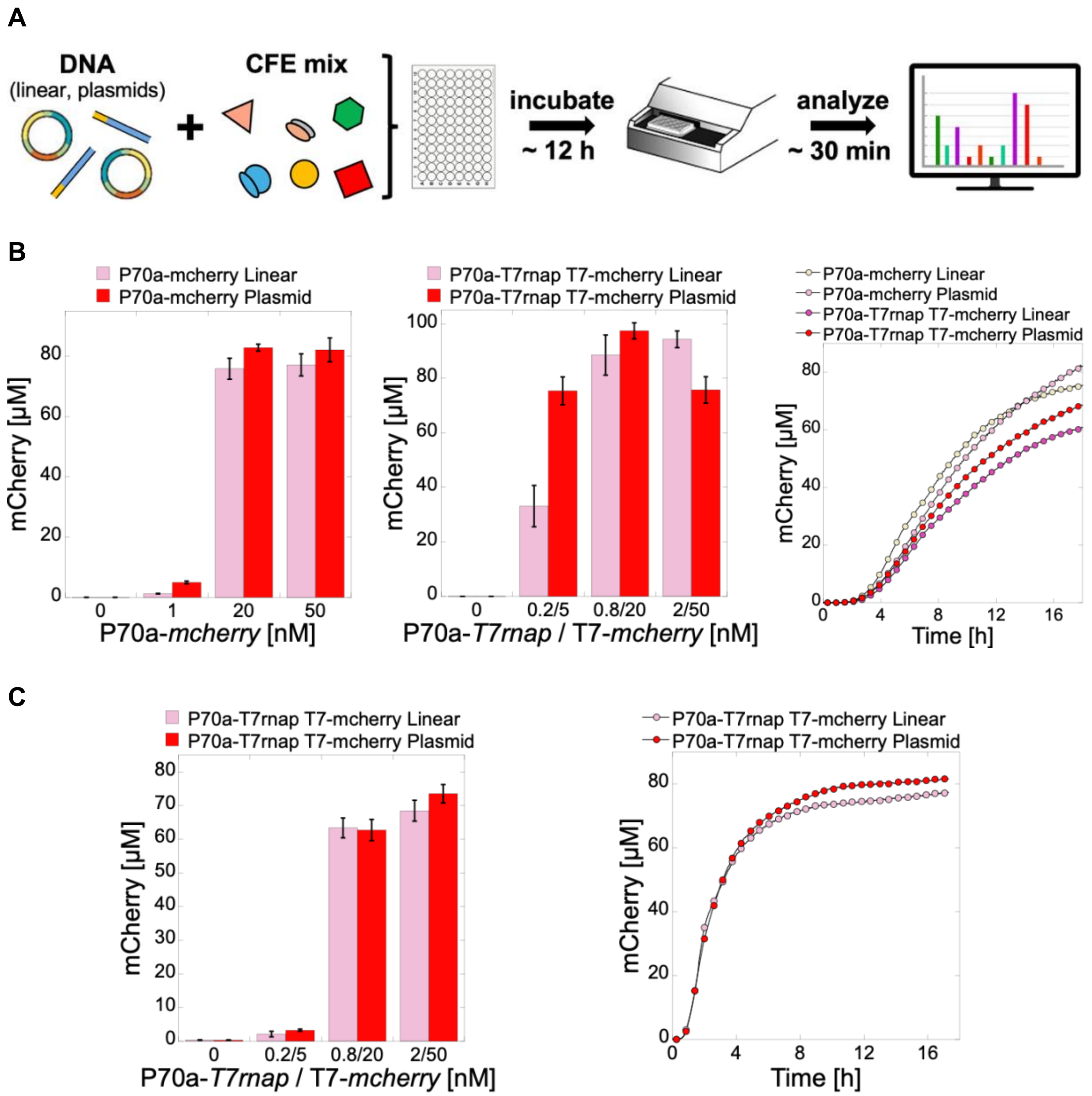


**Supplementary Figure 1.** CFE in batch mode. **A.** Diagram of the CFE workflow. **B.** Endpoint mCherry cell-free synthesis using the standard CFE system with linear DNA or plasmids, from the constitutive P70a promoter and the T7 transcriptional cascade. Kinetics of mCherry cell-free synthesis (P70a-*mcherry*: 20 nM, P70a-*T7rnap* 0.8 nM and T7-*mcherry* 20 nM). **C.** Endpoint mCherry cell-free synthesis using a ClearColi CFE system, using linear DNA or plasmids, from the T7 transcriptional cascade. Kinetics of mCherry cell-free synthesis (P70a-*T7rnap* 0.8 nM and T7-*mcherry* 20 nM).

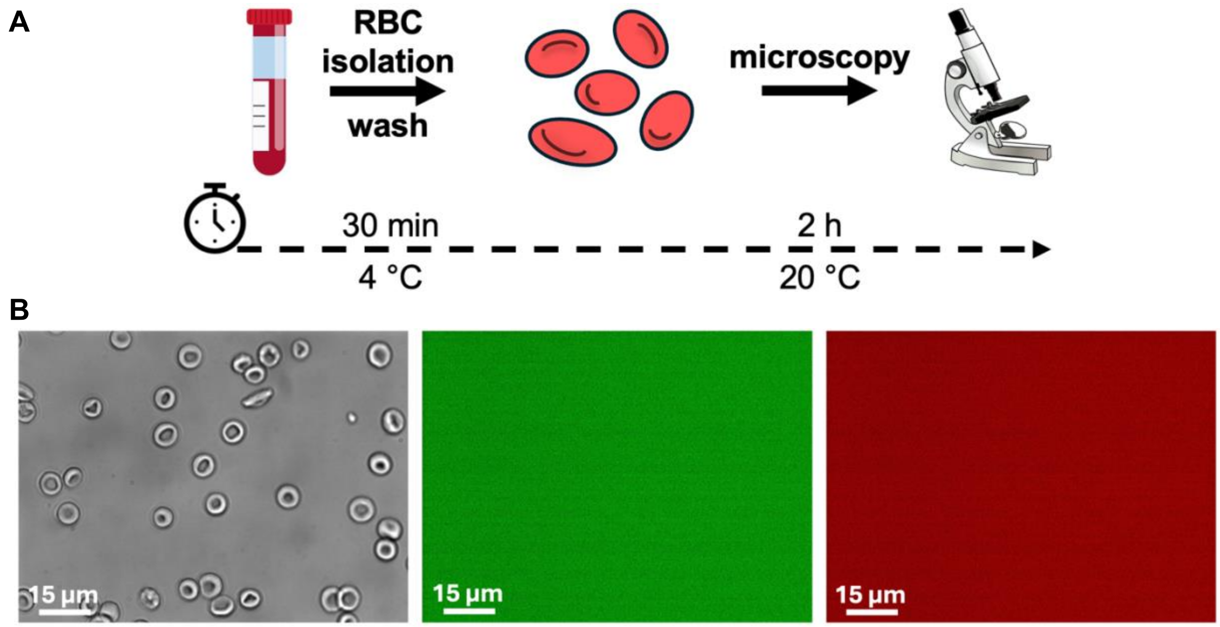


**Supplementary Figure 2.** Microscopy images of washed mouse RBCs. **A.** Diagram of the experiment. RBCs are isolated from mice, washed and observed by microscopy. RBCs were resuspended in buffer C1. **B.** Microscopy images of washed RBCs (phase contrasts, eGFP channel, mCherry channel).

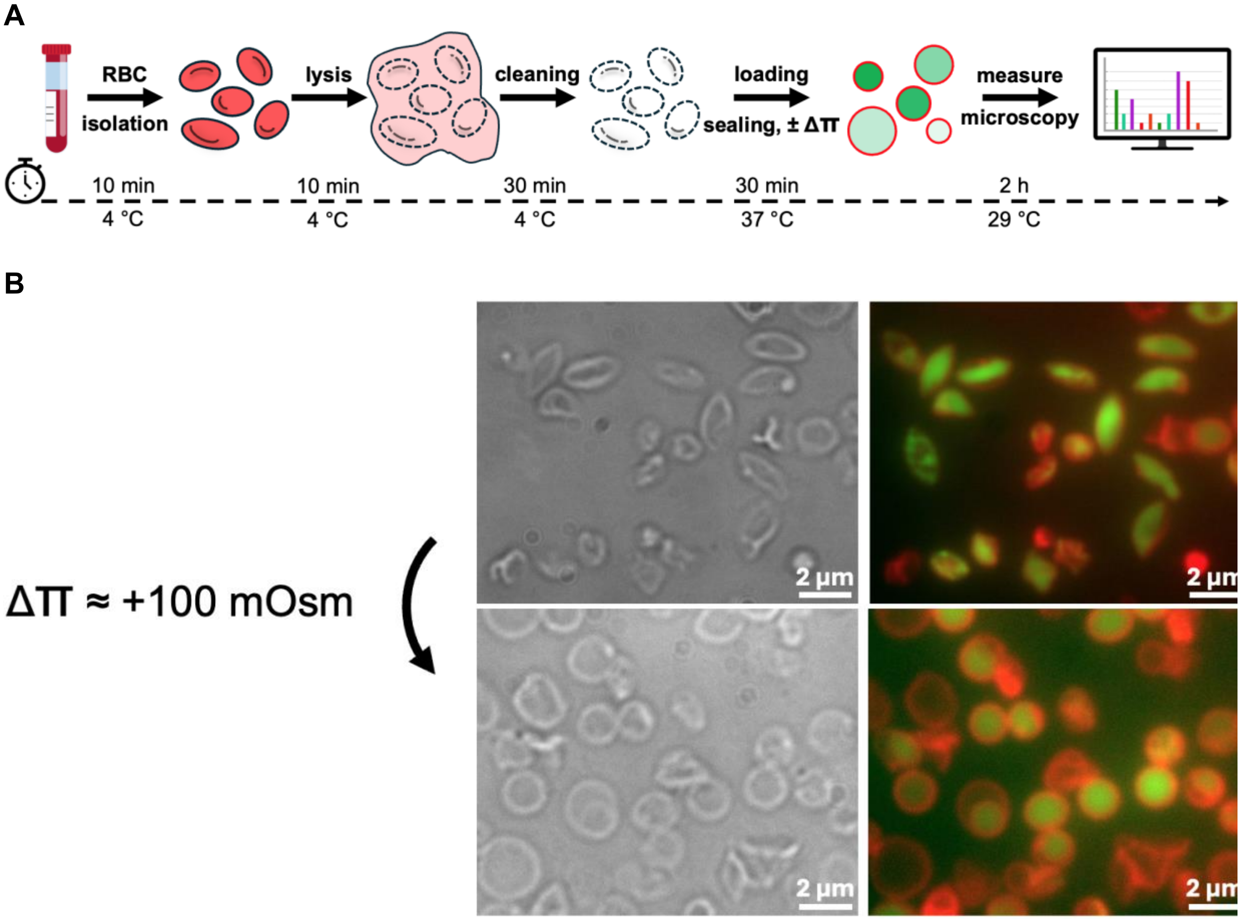


**Supplementary Figure 3.** Preparation and labeling of RBSCs. **A.** Diagram showing the steps of RBC ghosts' preparation and loading with or without osmotic stress. **B.** Microscopy images of RBC ghosts (phase contrast and merged of eGFP and mCherry channels). Top: iso-osmotic conditions, bottom: hypo-osmotic conditions that inflate RBSCs into spheres. RBC ghosts were loaded with FITC and the membrane labeled with Nile Red.

**A**

0 μM: $[A.U.] = -7800.7 + 16838 \times R^2$ $R = 0.85411$

1 μM: $[A.U.] = -2.8688\ 10^2 + 4.6222\ 10^5 \times R^2$ $R = 0.91661$

20 μM: $[A.U.] = -2.2767\ 10^2 + 7.0771\ 10^6 \times R^2$ $R = 0.97449$

50 μM: $[A.U.] = -3.4462\ 10^2 + 2.0659\ 10^2 \times R^2$ $R = 0.95277$

**B** **C**

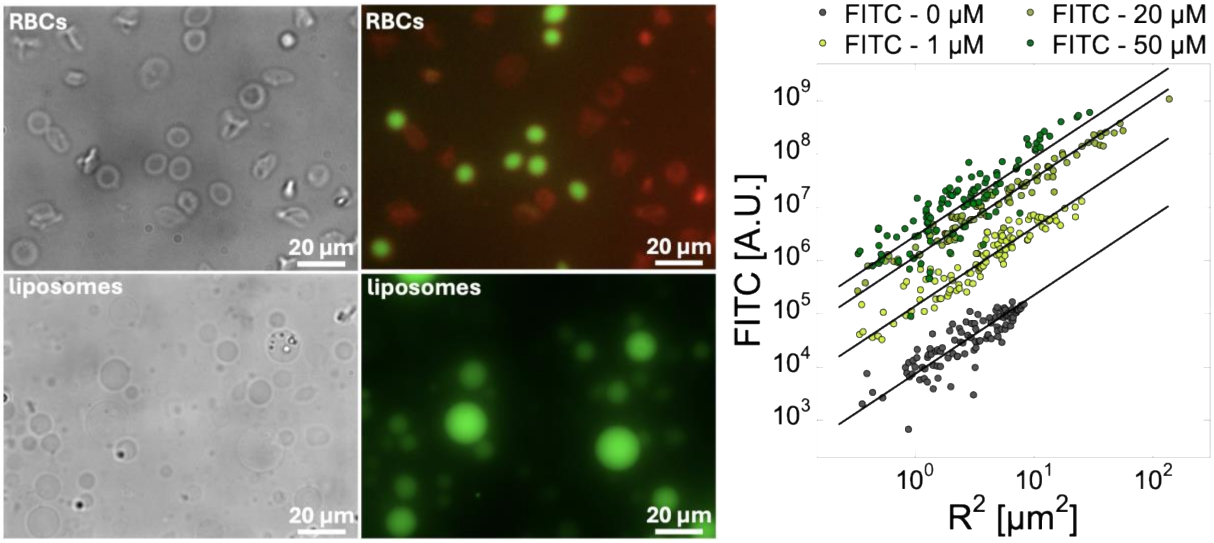


**Supplementary Figure 4.** Calibration of FITC fluorescence intensity into RBSCs. FITC was added to blank CFE reactions (no DNA) and encapsulated either into RBSCs or liposomes. **A.** Fit equations. **B.** Microscopy images of (top) RBC ghosts under osmotic stress to shape them into spheres, (bottom) liposomes. **C.** Graph of FITC intensity as a function of the liposomes' radii squared. This calibration was used to determine the concentration of FITC encapsulated into RBSCs.

**A**

0 μM: [A.U.] = -2433.7 + 3143.7 x $R^2$ R = 0.82002
1 μM: [A.U.] = -56408 + 45138 x $R^2$ R = 0.95611
20 μM: [A.U.] = -7.9222 $10^5$ + 8.4839 $10^5$ x $R^2$ R = 0.94993
50 μM: [A.U.] = -1.1717 $10^6$ + 1.5697 $10^6$ x $R^2$ R = 0.88589

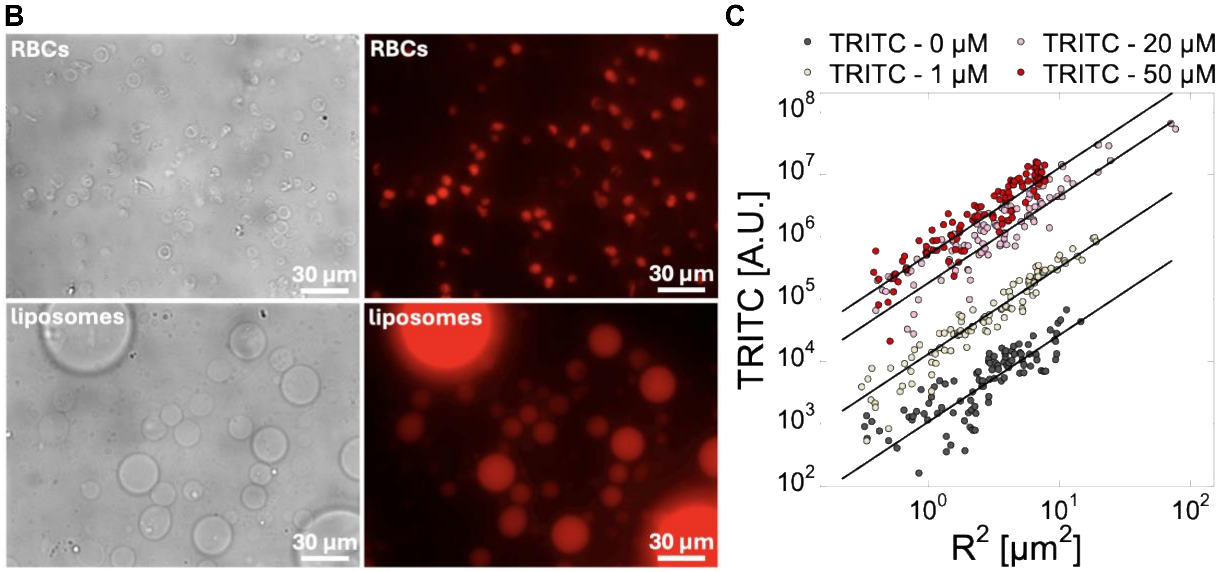


**Supplementary Figure 5.** Calibration of dextran-TRITC (40 kDa) fluorescence intensity into RBSCs. Dextran-TRITC was added to blank CFE reactions (no DNA) and encapsulated either into RBSCs or liposomes. **A.** Fit equations. **B.** Microscopy images of (top) RBC ghosts under osmotic stress to shape them into spheres, (bottom) liposomes. **C.** Graph of dextran-TRITC fluorescence intensity as a function of the liposomes' radii squared. This calibration was used to determine the concentration of dextran-TRITC encapsulated into RBSCs.

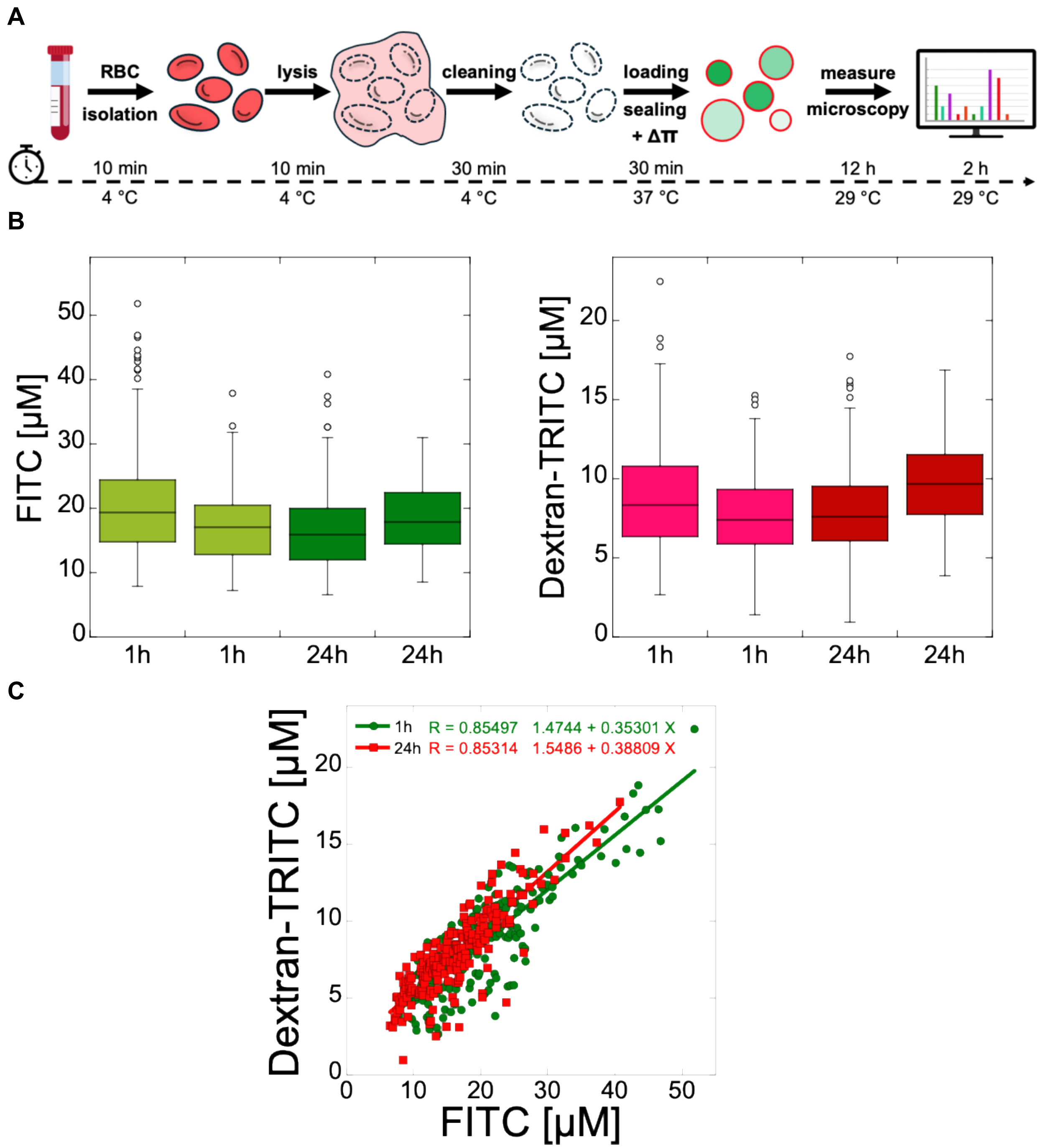


**Supplementary Figure 6.** Quantification of dyes encapsulation into RBSCs. **A.** Diagram of RBC ghosts' preparation. RBC ghosts were loaded with a blank CFE reaction (no DNA) containing either FITC (50 μM) or dextran-TRITC (40 kDa, 50 μM). **B.** The concentration of fluorescent dyes was determined using the calibrations. **C.** Both fluorescent dyes were encapsulated into RBSCs to determine the degree of encapsulation correlation. The data show a relatively high correlation. Two repeats were made, one measured after 1 hour, and the other one measured after 24 hours of incubation.

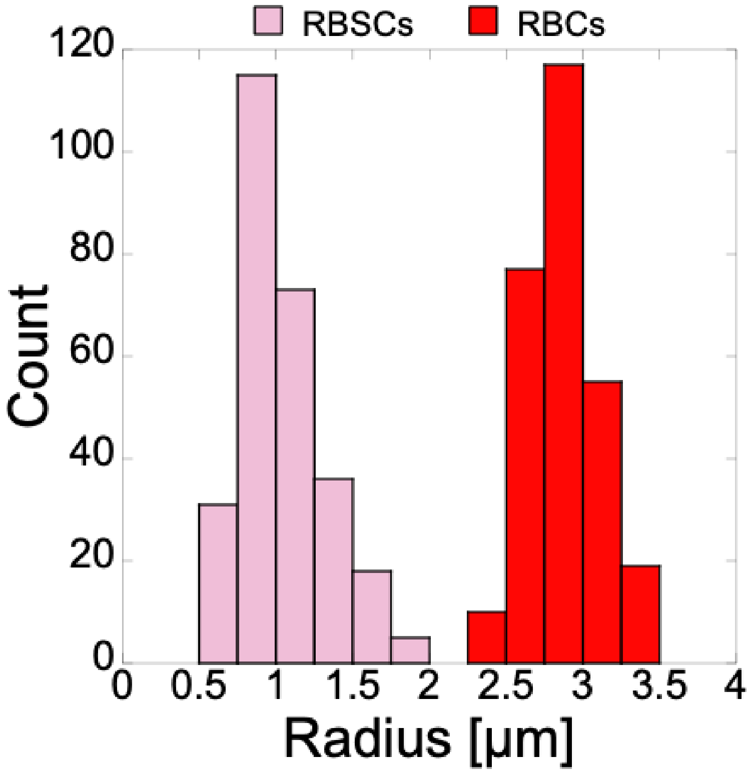


**Supplementary Figure 7.** Distribution of the RBCs and RBSCs radii. The radii of 250 RBCs or RBSCs were measured. RBCs and RBSCs were shaped into spheres under a slight osmotic stress to measure the radii.

**A**

0 µM: $[A.U.] = 9659 + 2577.6 \times R^2$ $\quad R = 0.97574$

1 µM: $[A.U.] = -18252 + 4916.4 \times R^2$ $\quad R = 0.94982$

20 µM: $[A.U.] = 1.0097\ 10^5 + 90279 \times R^2$ $\quad R = 0.97363$

50 µM: $[A.U.] = -4.3827\ 10^5 + 2.2043\ 10^5 \times R^2$ $\quad R = 0.955$

**B** **C**

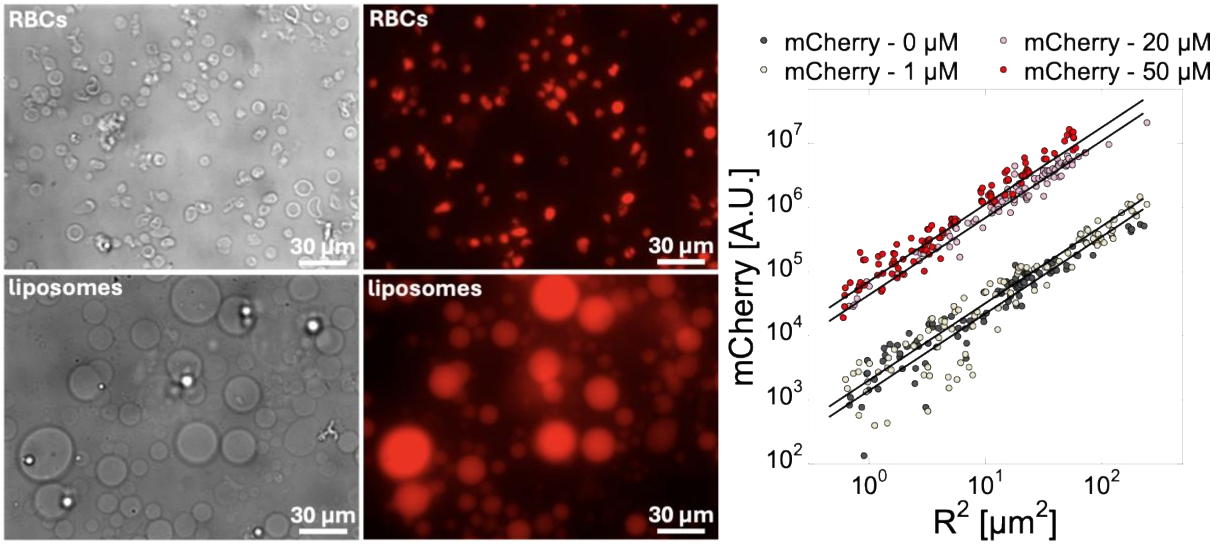


**Supplementary Figure 8.** Calibration of mCherry fluorescence intensity into RBSCs. Pure mCherry was added to blank CFE reactions (no DNA) and encapsulated into RBSCs. **A.** Fit equations. **B.** Microscopy images of (top) RBSCs under osmotic stress to shape them into spheres, (bottom) liposomes. **C.** Graph of mCherry fluorescence intensity as a function of the liposomes' radii squared. This calibration was used to determine the concentration of mCherry encapsulated into RBSCs.

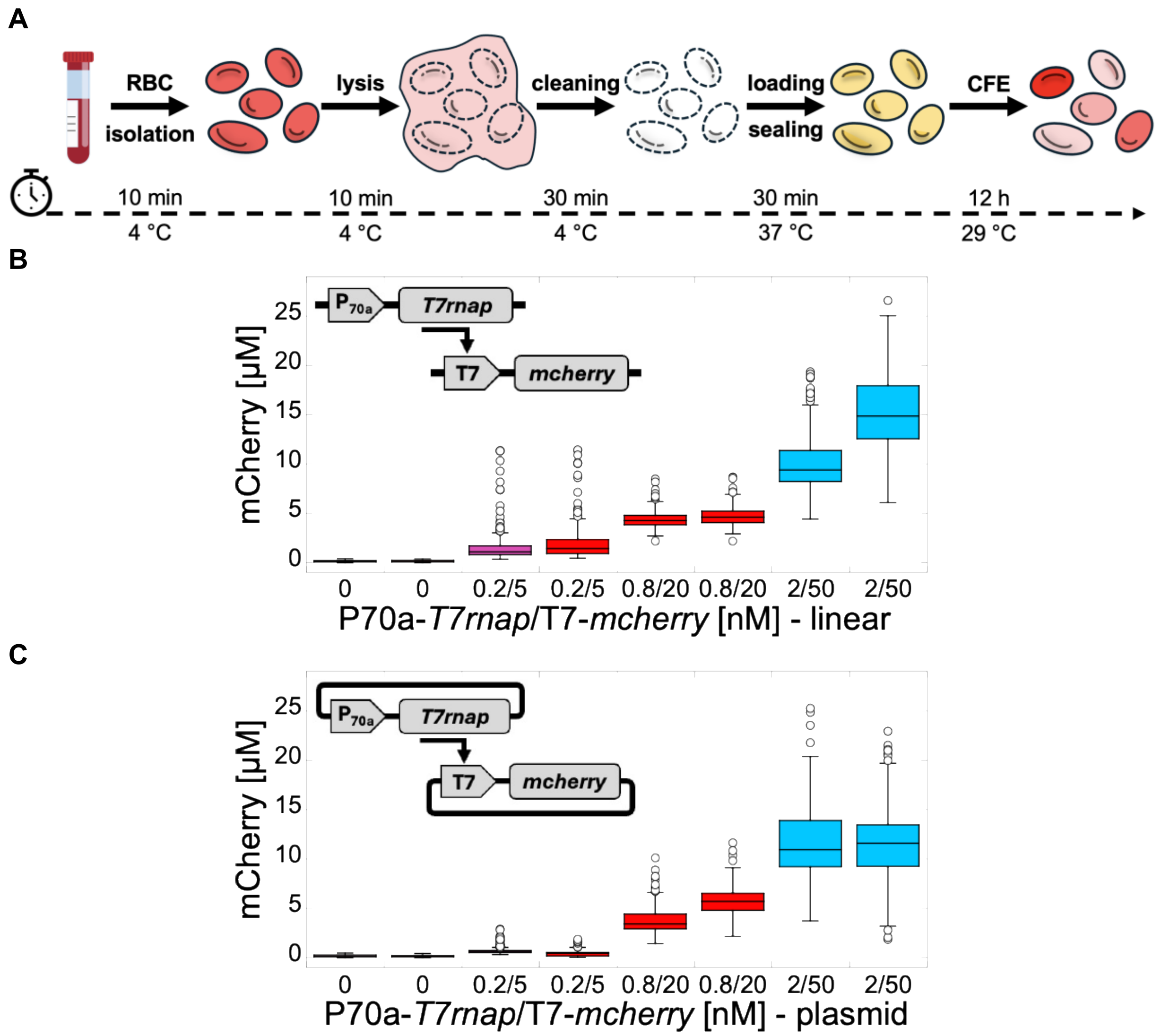


**Supplementary Figure 9.** CFE of *mcherry* into RBSCs using a ClearColi CFE system. **A.** Diagram of RBSCs preparation to express *mcherry* using a ClearColi CFE system. **B.** Box plot of mCherry cell-free synthesis through the T7 transcriptional activation cascade inside RBSCs using linear DNA, after 12 hours of incubation. Two repeats of each DNA concentration are shown. **C.** Box plot of mCherry cell-free synthesis through the T7 transcriptional activation cascade inside RBSCs using plasmids, after 12 hours of incubation. Two repeats of each DNA concentration are shown.

**A**

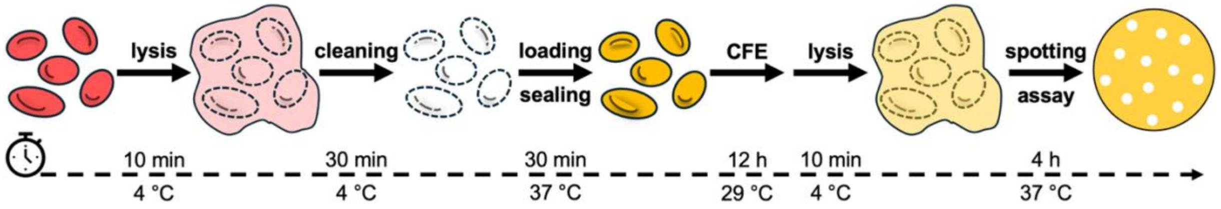


**B**

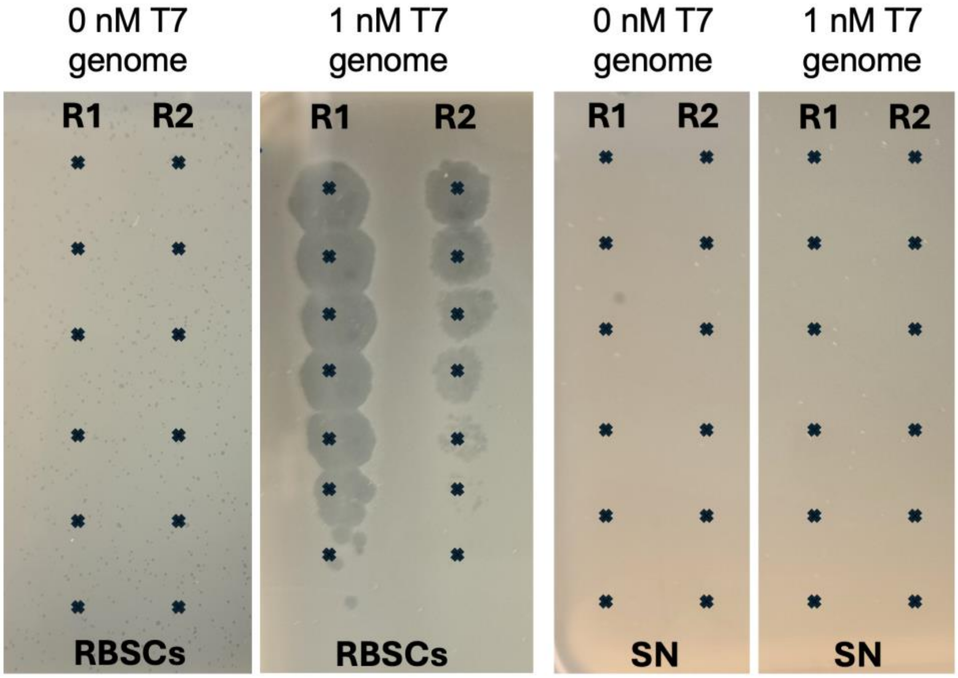


**C**

| | RBC | Supernatant |
|---|---|---|
| 0 nM T7 genome | 0 PFU/ml | 0 PFU/ml |
| 0 nM T7 genome | 0 PFU/ml | 0 PFU/ml |
| 1 nM T7 genome | 6.26 $10^{10}$ PFU/ml | 0 PFU/ml |
| 1 nM T7 genome | 4.67 $10^{9}$ PFU/ml | 0 PFU/ml |

**Supplementary Figure 10.** Cell-free synthesis of phage T7 inside RBSCs. **A.** Diagram of phage T7 cell-free synthesis into RBSCs. **B.** Spotting assay of T7 cell-free synthesis (two repeats) at two T7 genome concentrations into RBSCs. No phages were found in the supernatant (SN). **C.** Table reporting the plaque forming units per milliliter (PFU/ml) for the two repeats.

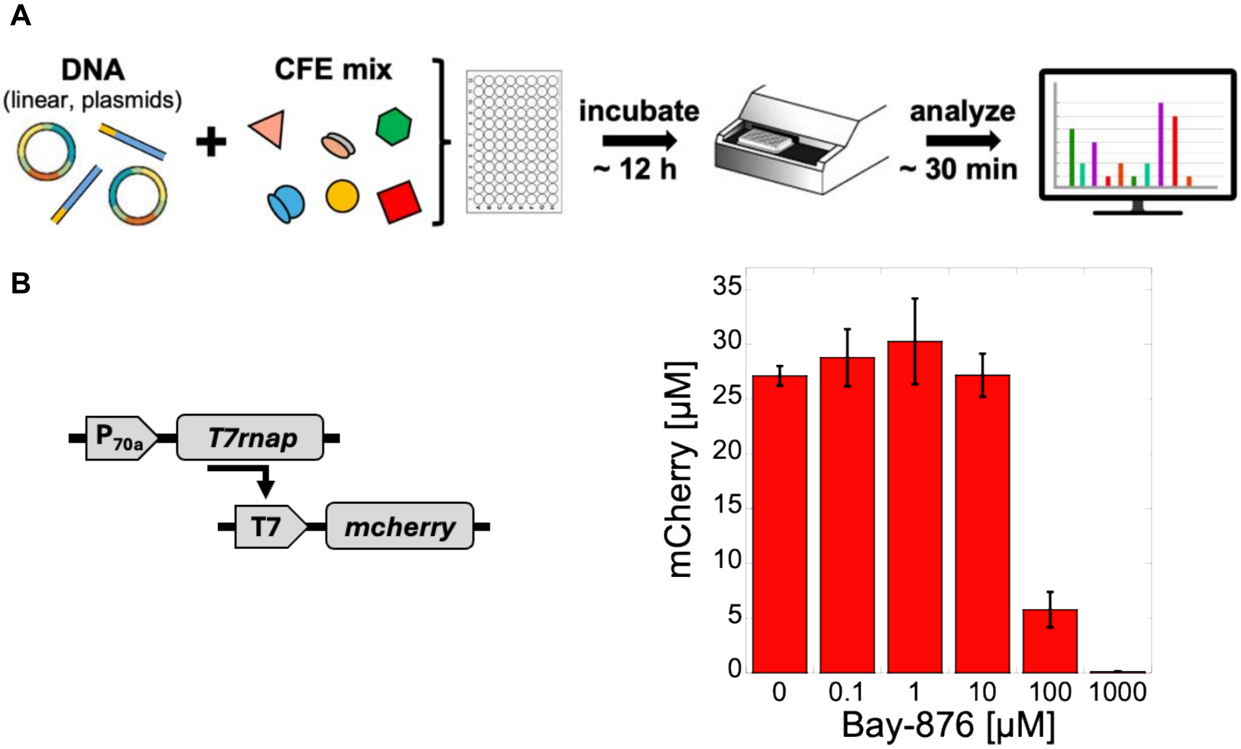


**Supplementary Figure 11.** Effect of Bay-876 on CFE. **A.** Workflow of the experiment carried out in batch mode. **B.** mCherry was synthesized from a T7 transcriptional activation cascade using linear DNA (P70a-*T7rnap* 0.2 nM, T7-*mcherry* 10 nM). **C.** Endpoint mCherry cell-free synthesis as a function of Bay-876 added to the reactions (measured after 12 hours of incubation).

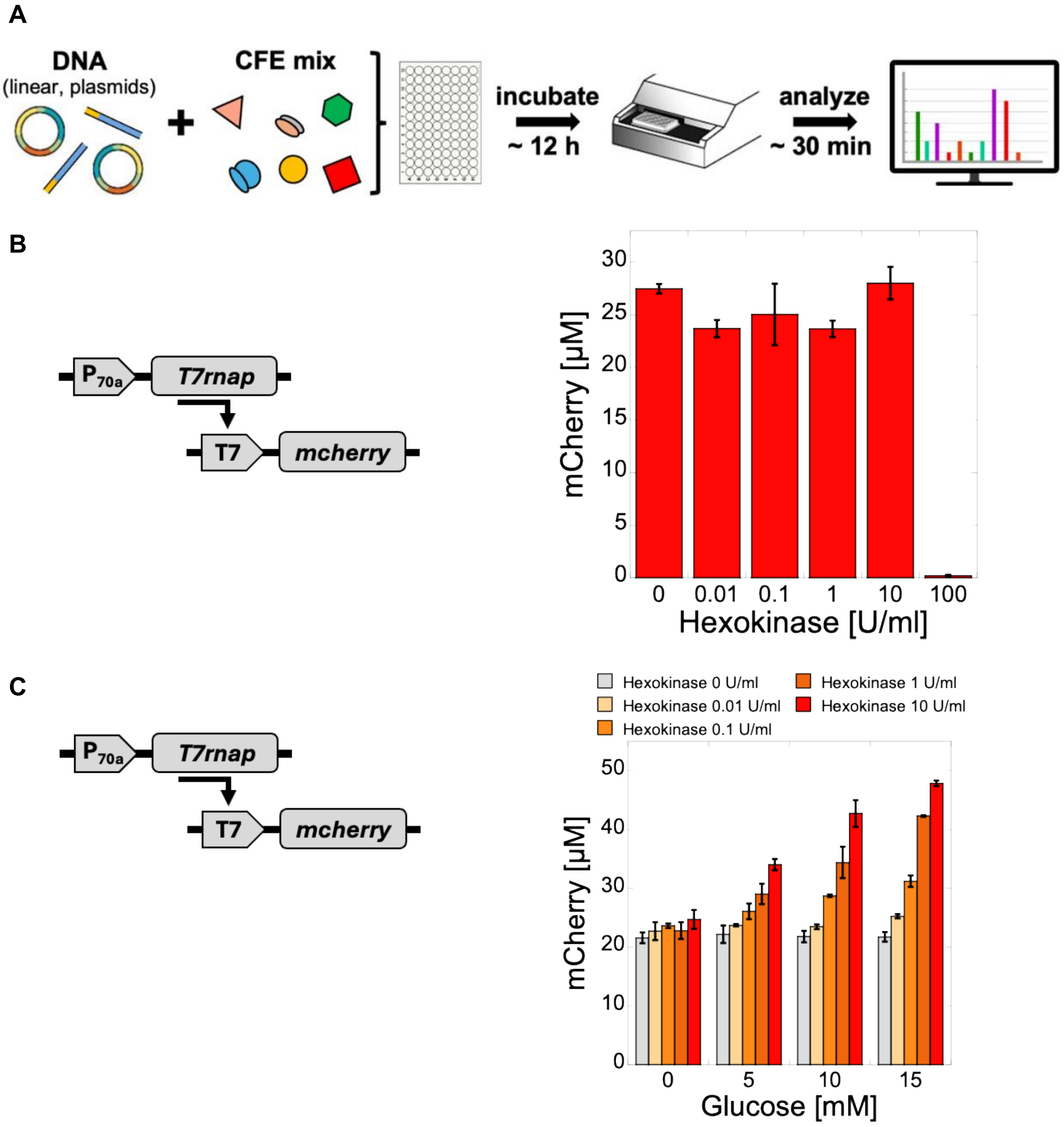


**Supplementary Figure 12.** Effect of hexokinase on CFE. **A.** Workflow of the experiment. **B.** mCherry was synthesized from a T7 transcriptional activation cascade using linear DNA (P70a-*T7rnap* 0.2 nM, T7-*mcherry* 10 nM). Endpoint mCherry cell-free synthesis as a function of hexokinase added to the reactions. **C.** Endpoint mCherry cell-free synthesis as a function of hexokinase and glucose added to the reactions, using linear DNA (measured after 12 hours of incubation).

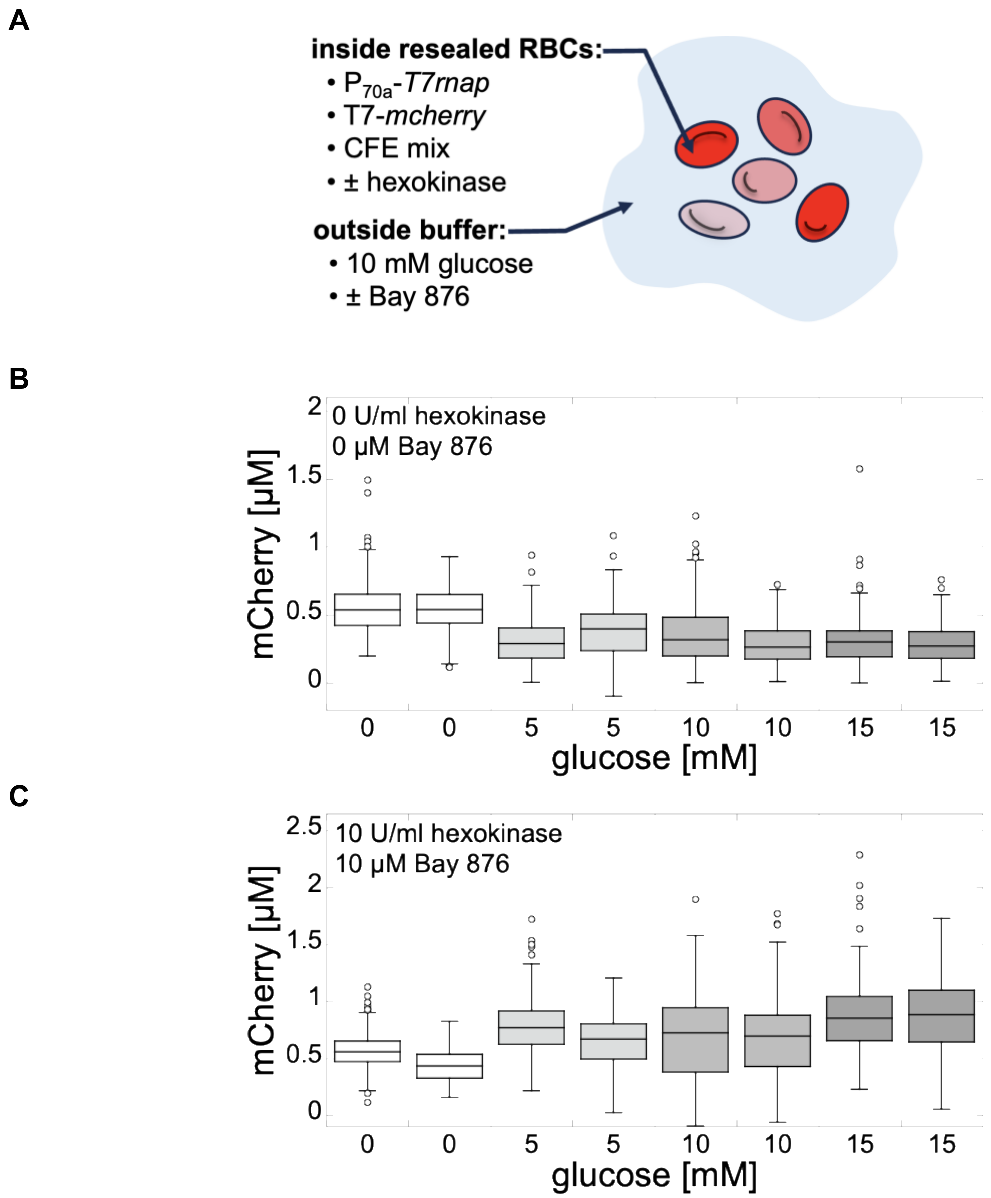


**Supplementary Figure 13.** A glucose sensor system. **A.** Diagram of the system using RBSCs. *mcherry* was expressed through the T7 promoter (transcriptional activation cascade, using plasmids). **B.** Box plots of mCherry cell-free synthesis inside RBSCs using plasmids, without hexokinase and without Bay-876. Two repeats of each glucose concentration are shown. The fluorescence of two hundred RBSCs was measured and quantified after 12 hours of incubation for each glucose concentration. **C.** Box plots of mCherry cell-free synthesis inside RBSCs using plasmids, with hexokinase and with Bay-876. Two repeats of each glucose concentration are shown. The fluorescence of two hundred RBSCs was measured and quantified after 12 hours of incubation for each glucose concentration.

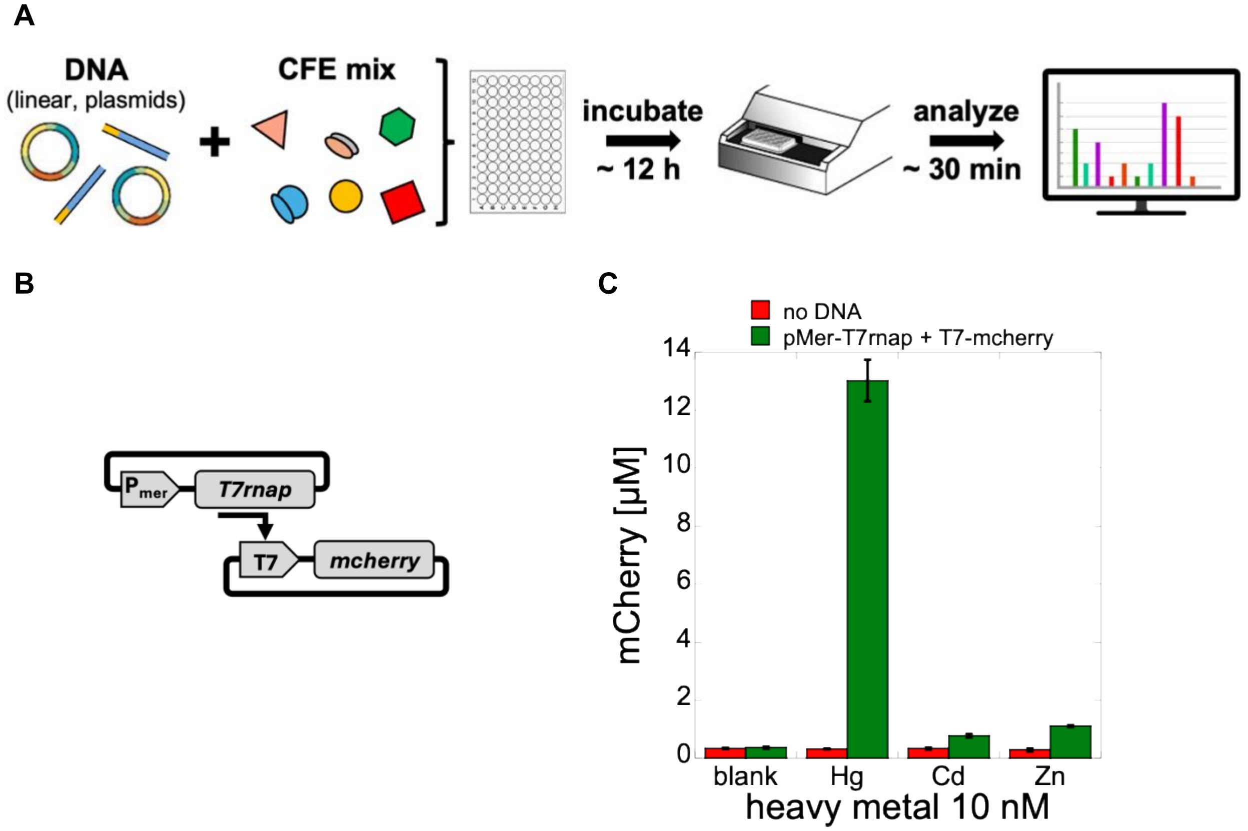


**Supplementary Figure 14.** CFE using the Pmer inducible system in batch mode CFE reactions. **A.** Workflow of the experiment. **B.** mCherry was synthesized from a T7 transcriptional activation cascade using plasmids (Pmer-*T7rnap* at 0.2 nM, T7-*mcherry* at 5 nM). **C.** Endpoint mCherry cell-free synthesis from the Pmer transcriptional activation cascade as a function of 10 nM heavy metal added to the reaction, measured after 12 hours of incubation. Blank: no heavy metal added to the CFE reaction.

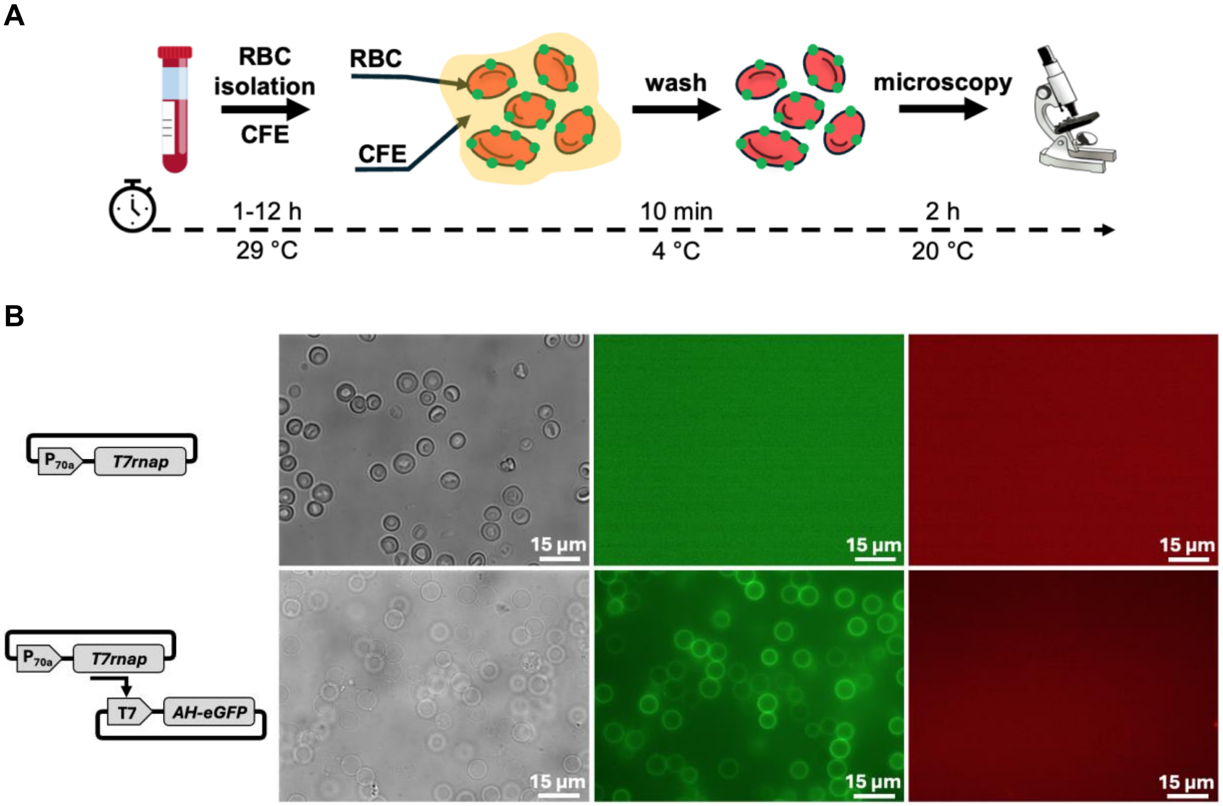


**Supplementary Figure 15.** Cell-free synthesis of alpha-hemolysin (AH) in the presence of RBCs. **A.** Workflow of the experiment. RBCs were added to the cell-free reaction. **B.** Microscopy images of RBCs after CFE of either *T7rnap* or *AH-eGFP* (plasmids: P70a-*T7rnap* 0.2 nM, T7-*AH-eGFP* 4 nM). Microscopy channels from left to right: phase contrast, eGFP, mCherry. Images were taken after 2 hours of incubation.

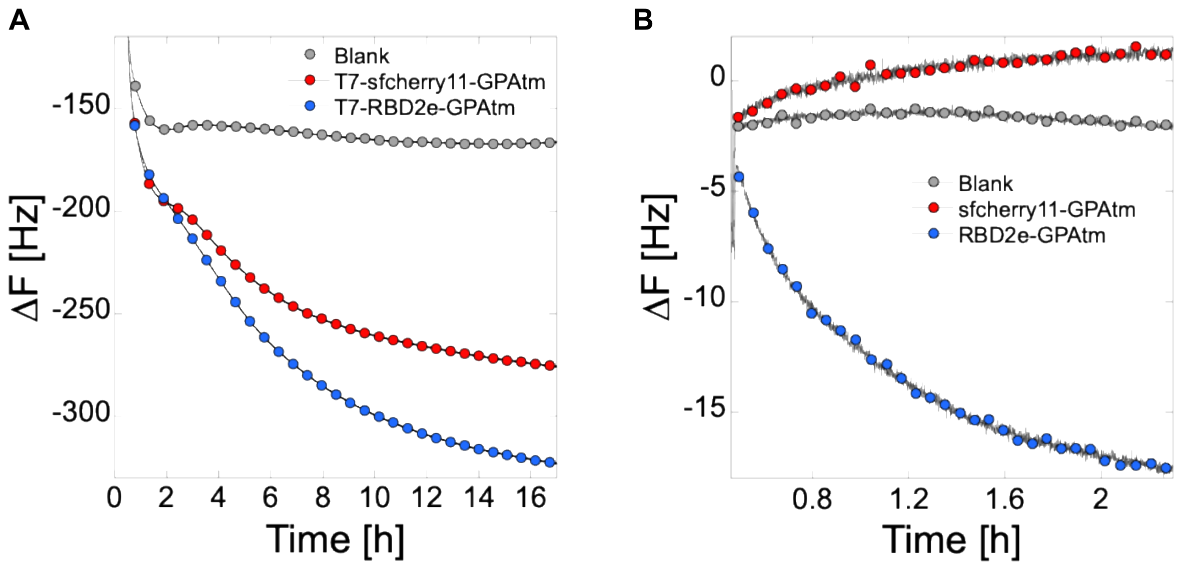


**Supplementary Figure 16.** Anchoring payloads to phospholipid bilayers using GPAtm (QCMD). **A.** Cell-free synthesized sfcherry11-GPAtm (P70a-*T7rnap* 2 nM, T7-*sfcherry11-GPAtm* 50 nM) and RBD2e-GPAtm (P70a-*T7rnap* 2 nM, T7-*RBD2e-GPAtm* 50 nM) are both inserted into an *E. coli* (ECL) supported lipid bilayers (SLBs) as indicated by the drop in frequency shift. Blank reaction consisted of a CFE reaction without added DNA. **B.** The antibody 5G8 (0.5 μM) specific to RBD2e was incubated on three different ECL SLBs: a blank SLB, an SLB after cell-free synthesis of sfcherry11-GPAtm and an SLB after cell-free synthesis of RBD2e-GPAtm. 5G8 only binds to the SLB having RBD2e-GPAtm, as noticed by the drop in frequency shift.

| | LPS [U/ml] |
|---|---|
| Lysate 1 | 36700 ± 23100 |
| Lysate 2 | 26667 ± 20817 |
| Lysate 3 | 36667 ± 15275 |
| ClearColi lysate | 1 ± 0 |

**Supplementary Table 1.** Concentration of LPS measured in three different *E. coli* lysates and a ClearColi lysate.

| | LPS [U/ml] |
|---|---|
| CFE reaction with lysate 1 + RBCs | 0 |
| CFE reaction with lysate 2 + RBCs | 0 |
| CFE reaction with lysate 3 + RBCs | 0 |
| CFE reaction with ClearColi lysate + RBCs | 0 |

**Supplementary Table 2.** Concentration of LPS measured after washing RBCs incubated with a CFE reaction. The RBCs were isolated from blood, washed and incubated in four different types of CFE reactions (based on three different *E. coli* lysate batches and one ClearColi lysate). mCherry was cell-free synthesized (plasmid P70a-*mcherry* 10 nM). The RBCs were subsequently washed after 12 hours of incubation and resuspended in a buffer. The concentration of LPS was measured.
